# Generation of twenty four induced pluripotent stem cell lines from twenty four members of the Lothian Birth Cohort 1936

**DOI:** 10.1101/2020.02.05.935213

**Authors:** Jamie Toombs, Lindsay Panther, Loren Ornelas, Chunyan Liu, Emilda Gomez, Raquel Martín-Ibáñez, Simon R. Cox, Stuart J. Ritchie, Sarah E. Harris, Adele Taylor, Paul Redmond, Tom C. Russ, Lee Murphy, James D. Cooper, Karen Burr, Bhuvaneish T. Selvaraj, Cathy Browne, Clive N. Svendsen, Sally A. Cowley, Ian J. Deary, Siddharthan Chandran, Tara Spires-Jones, Dhruv Sareen

## Abstract

Cognitive decline is among the most feared aspects of ageing. We have generated induced pluripotent stem cells (iPSCs) from 24 people from the Lothian Birth Cohort 1936, whose cognitive ability was tested in childhood and in older age. Peripheral blood mononuclear cells (PBMCs) were reprogrammed using non-integrating oriP/EBNA1 backbone plasmids expressing six iPSC reprogramming factors (OCT3/4 (POU5F1), SOX2, KLF4, L-Myc, shp53, Lin28, SV40LT). All lines demonstrated STR matched karyotype and pluripotency was validated by multiple methods. These iPSC lines are a valuable resource to study molecular mechanisms underlying individual differences in cognitive ageing and resilience to age-related neurodegenerative diseases.

## Resource Table

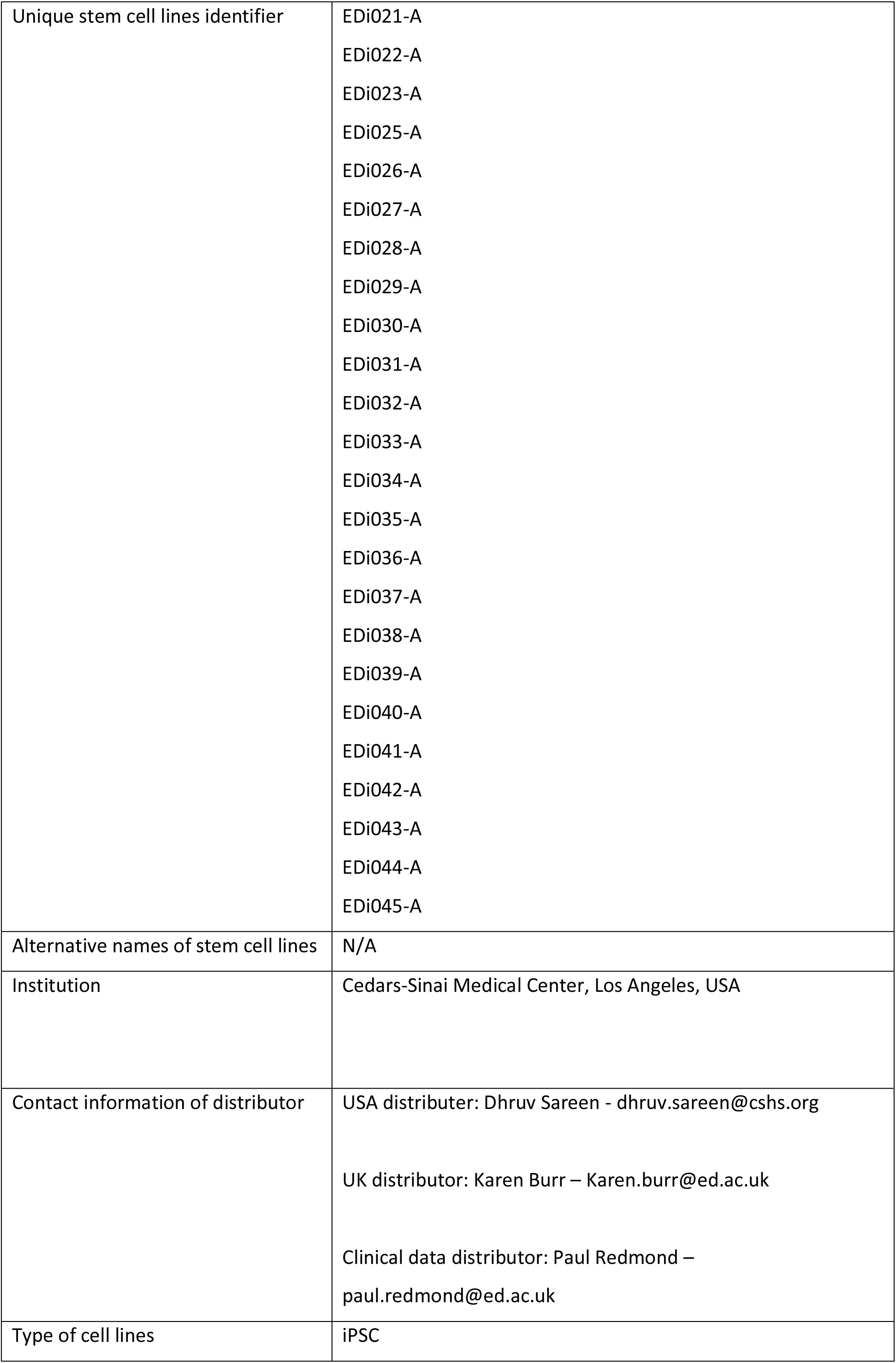
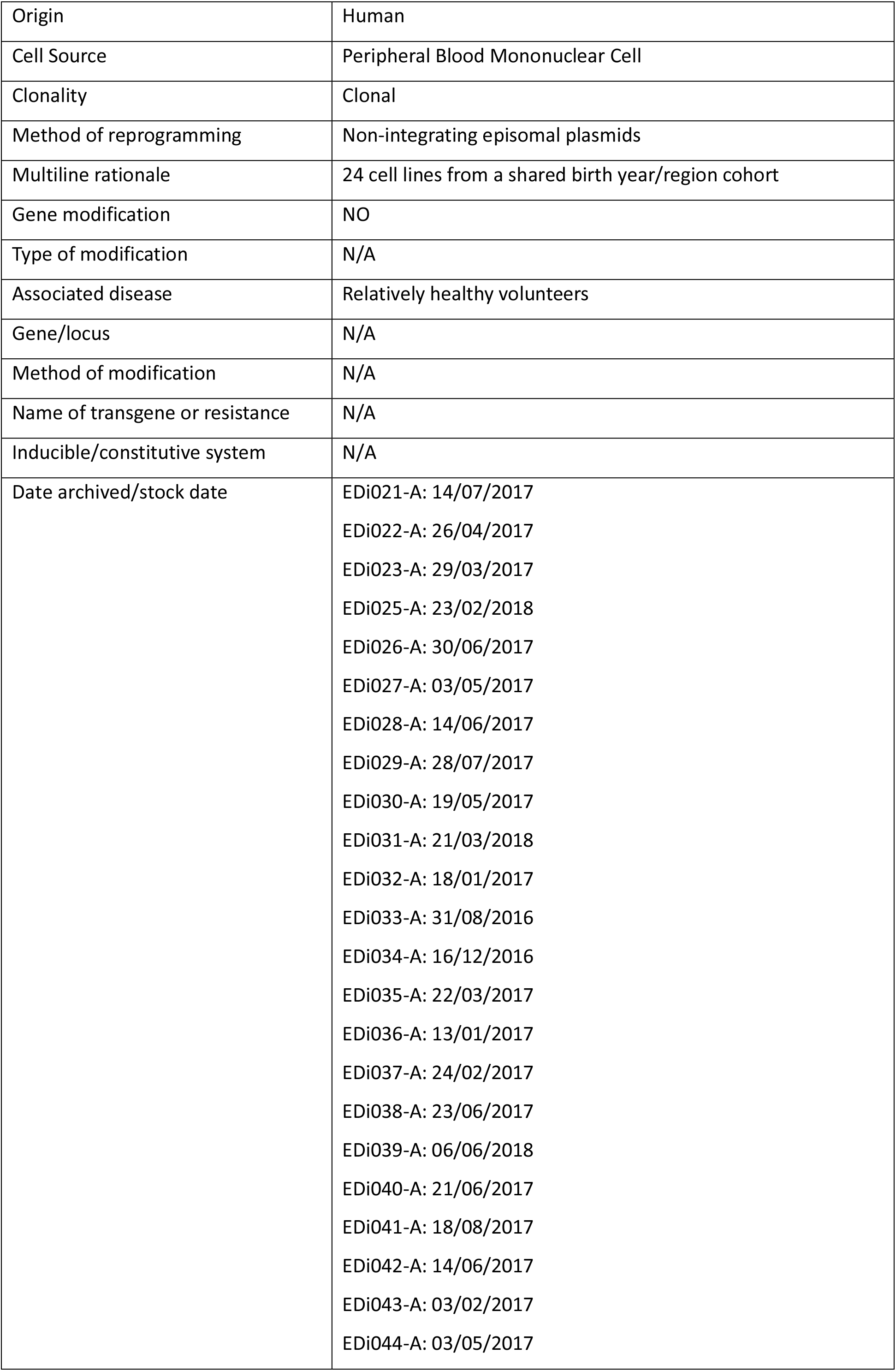
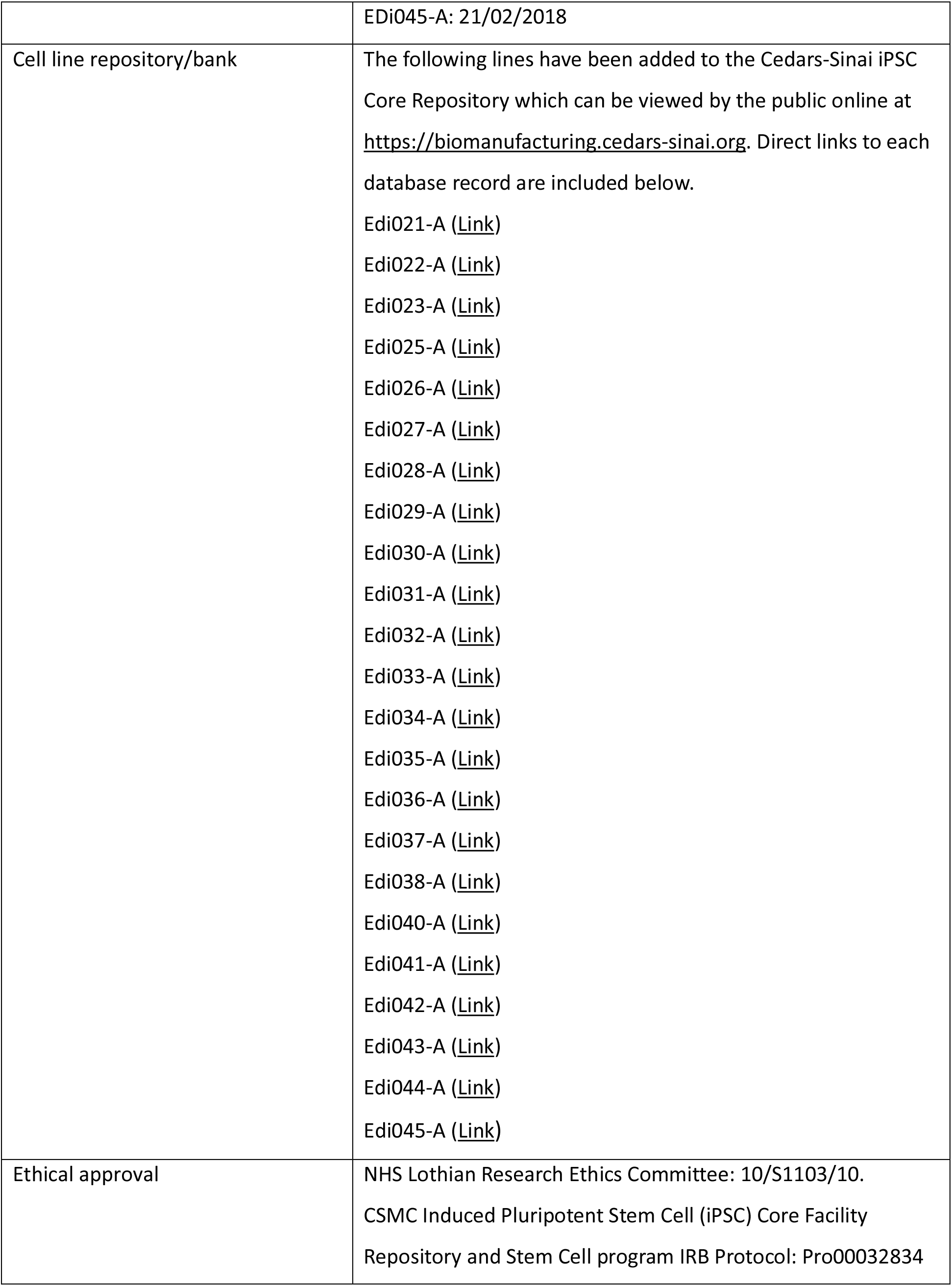

## Resource utility

The neurobiology of cognitive ability and its decline during ageing are poorly understood. Human iPSC lines from the Lothian Birth Cohort 1936 comprise individuals with rich life-course cognitive performance data (Taylor et al., 2018; Wardlaw et al., 2011), affording a rare model to investigate molecular mechanisms relevant to differences in brain development, cellular resilience, and vulnerability to pathology.

## Resource Details

Human peripheral blood mononuclear cells (PBMCs) were obtained from 24 unrelated members of the Lothian Birth Cohort 1936. Demographic parameters are 50% female (n = 12), 100% white Scottish (Table 1). Line donors can be grouped into ‘successful’, ‘typical’, and ‘poor’ cognitive ageing categories (sFig.1). Exclusion criteria were: self-reported dementia, Parkinson’s disease or stroke, Mini Mental State Examination (MMSE score <24, as well as standardised childhood IQ scores (<65, Moray House Test No. 12 at age 11), and standardised adult IQ scores (<85, average of Moray House Test No. 12 at age 70 and 76).

**Table 1:** Summary of lines

| **iPSC line names** | **Abbreviation in figures** | **Gender** | **Age at collection** | **Ethnicity** | **Genotype of locus** | **Disease** |
| --- | --- | --- | --- | --- | --- | --- |
| EDi021-A |  | M | 78.8 | White<br>Scottish | N/A | N/A |
| EDi022-A |  | M | 79.22 | White<br>Scottish | N/A | N/A |
| EDi023-A |  | F | 79.1 | White<br>Scottish | N/A | N/A |
| EDi025-A |  | M | 78 | White<br>Scottish | N/A | N/A |
| EDi026-A |  | M | 79.45 | White<br>Scottish | N/A | N/A |
| EDi027-A |  | F | 79.65 | White<br>Scottish | N/A | N/A |
| EDi028-A |  | M | 79.1 | White<br>Scottish | N/A | N/A |
| EDi029-A |  | M | 80.13 | White<br>Scottish | N/A | N/A |
| EDi030-A |  | F | 78.98 | White<br>Scottish | N/A | N/A |
| EDi031-A |  | F | 78 | White<br>Scottish | N/A | N/A |
| EDi032-A |  | F | 79.29 | White<br>Scottish | N/A | N/A |
| EDi033-A |  | F | 78.67 | White<br>Scottish | N/A | N/A |
| EDi034-A |  | F | 78.68 | White<br>Scottish | N/A | N/A |
| EDi035-A |  | F | 78.79 | White<br>Scottish | N/A | N/A |
| EDi036-A |  | F | 79.22 | White<br>Scottish | N/A | N/A |
| EDi037-A |  | M | 79.1 | White<br>Scottish | N/A | N/A |
| EDi038-A |  | M | 79.19 | White<br>Scottish | N/A | N/A |
| EDi039-A |  | M | 78 | White<br>Scottish | N/A | N/A |
| EDi040-A |  | M | 79.67 | White<br>Scottish | N/A | N/A |
| EDi041-A |  | F | 80.13 | White<br>Scottish | N/A | N/A |
| EDi042-A |  | F | 79.42 | White<br>Scottish | N/A | N/A |
| EDi043-A |  | M | 80.26 | White<br>Scottish | N/A | N/A |
| EDi044-A |  | F | 79.85 | White<br>Scottish | N/A | N/A |
| EDi045-A |  | M | 80.32 | White<br>Scottish | N/A | N/A |

PBMCs were reprogrammed to generate induced pluripotent stem cells (iPSCs) using episomal plasmids encoding human OCT3/4 (POU5F1), SOX2, KLF4, L-Myc, shp53, Lin28, SV40LT. All lines were reprogrammed and stored within 22 months of each other. EBNA-related gene analysis demonstrated that iPSCs were EBNA transgene-free (and therefore exogenous reprogramming factors were no longer present) by passage 17-21 (depending on line). Qualitative tests for parental cell type by TCR-αβ and TCR-γδ T-cell clonality assay revealed that 83% (n = 20) of lines were non-T cell-derived, 17% (n = 4) were T-cell derived. T-cell derived lines are: EDi021-A, EDi025-A, EDi026-A, and EDi035-A.

Pluripotency was confirmed by the expression of six pluripotency markers (OCT3/4, NANOG, SOX2, TRA-1-60, TRA-1-81, SSEA4) evaluated by immunocytochemistry (Fig.1B, sFig.2-24B). Additionally, all lines demonstrated positive alkaline phosphatase AP staining (Fig.1A, sFig.2-24A) and self-renewal in undifferentiated iPSCs as assessed by PluriTest (Fig.1C, sFig.2-24C) and TaqMan®hPSC Scorecard^TM^ Panel (Fig.1D, sFig.2-24E). However, whilst EDi035-A had a positive PluriTest and Scorecard^TM^ pluripotency result, the PluriTest novelty score was borderline (1.688) (sFig.14C,E). Furthermore, EDi027-A also had a borderline positive ectoderm score as assessed by Scorecard^TM^ (sFig.6E). At 14 days of embryoid body differentiation, all lines demonstrated tri-lineage potential except EDi022-A (negative endoderm, borderline mesoderm score, sFig.2E), EDi035-A (negative mesoderm, borderline endoderm score, sFig.14E), and EDi042-A (negative endoderm score, sFig.21E), as assessed by Scorecard^TM^.

**Figure 1:**
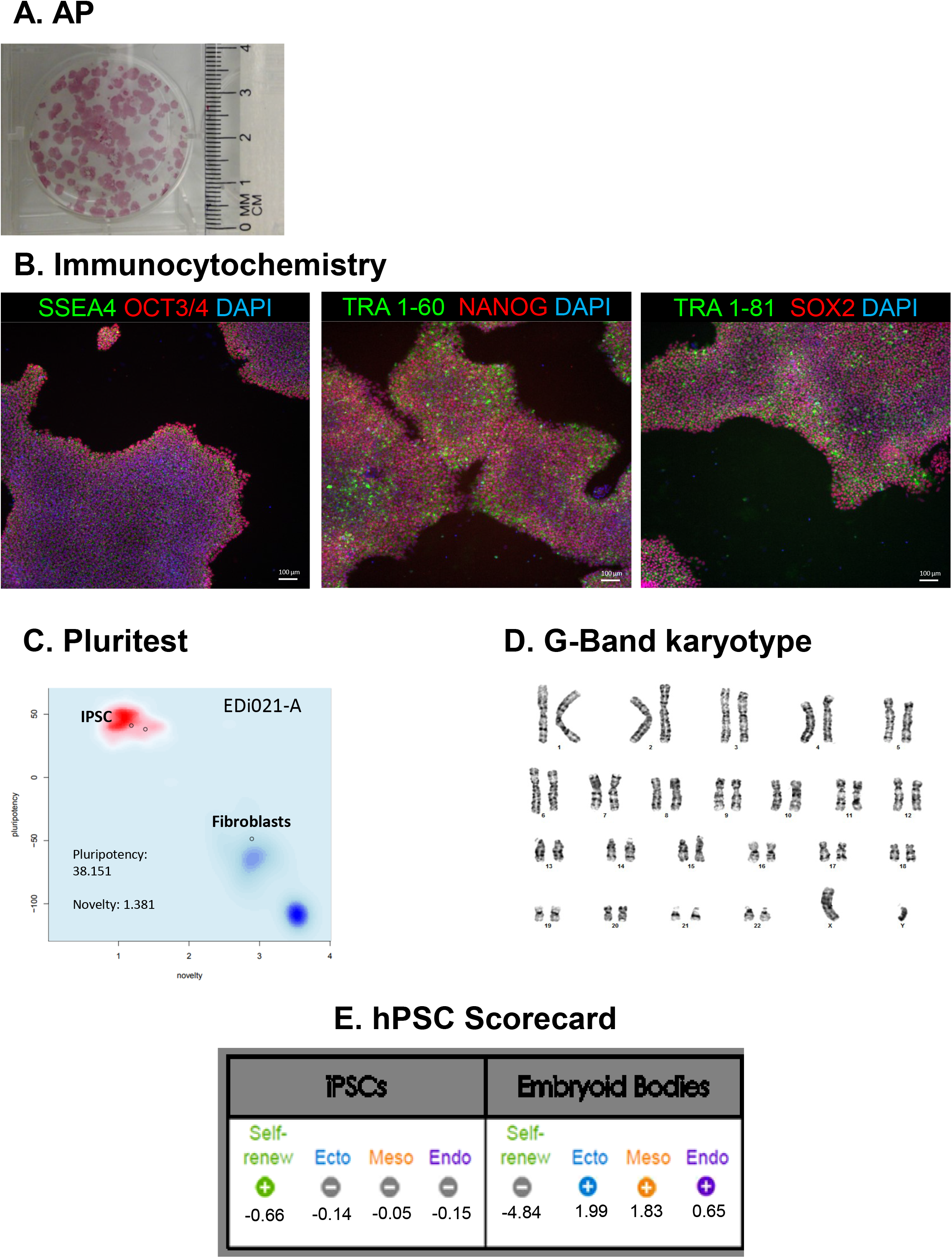
Characterization for iPSC line EDi021-A

All lines showed a normal karyotype (Fig.1D, sFig. 2-24D) between passages 6-22, with one exception. All five clones of EDi-038-A (a male) karyotyped as monosomy (45,X) (sFig.18D), and thus very likely stems from the source PBMCs. Mosaicism is a relatively common and probably harmless finding in blood cultures from normal females and, though rarer, also in males (Bukvic et al., 2001). No differences were detected between the original PBMC samples and the corresponding iPSC lines.

All lines were confirmed to be of human origin and iPSCs matched the profile of parent PBMCs by Short Tandem Repeat (STR) analysis. Parent line data was not available for EDi026-A and EDi028-A. Genetic profiles for these lines were compared to the cell line genetic profiles available in the DSMZ STR database and did not match any other reported profiles in the DSMZ database. These profiles were found to be unique and did not match to any previously submitted profiles from the iPSC Core. The genetic profiles established here can be used for future comparisons for these cell lines. Whole genome sequence data for all 24 lines has been deposited at the European Genome-phenome Archive (EGA), which is hosted by the EBI and the CRG, under accession number EGAS00001003819.

An overview of iPSC line characterisation can be found in Table 2. Figure 1 presents example characterisation data from EDi021-A. Data for all other lines can be found in Supplementary Figures 2-24.

**Table 2:**
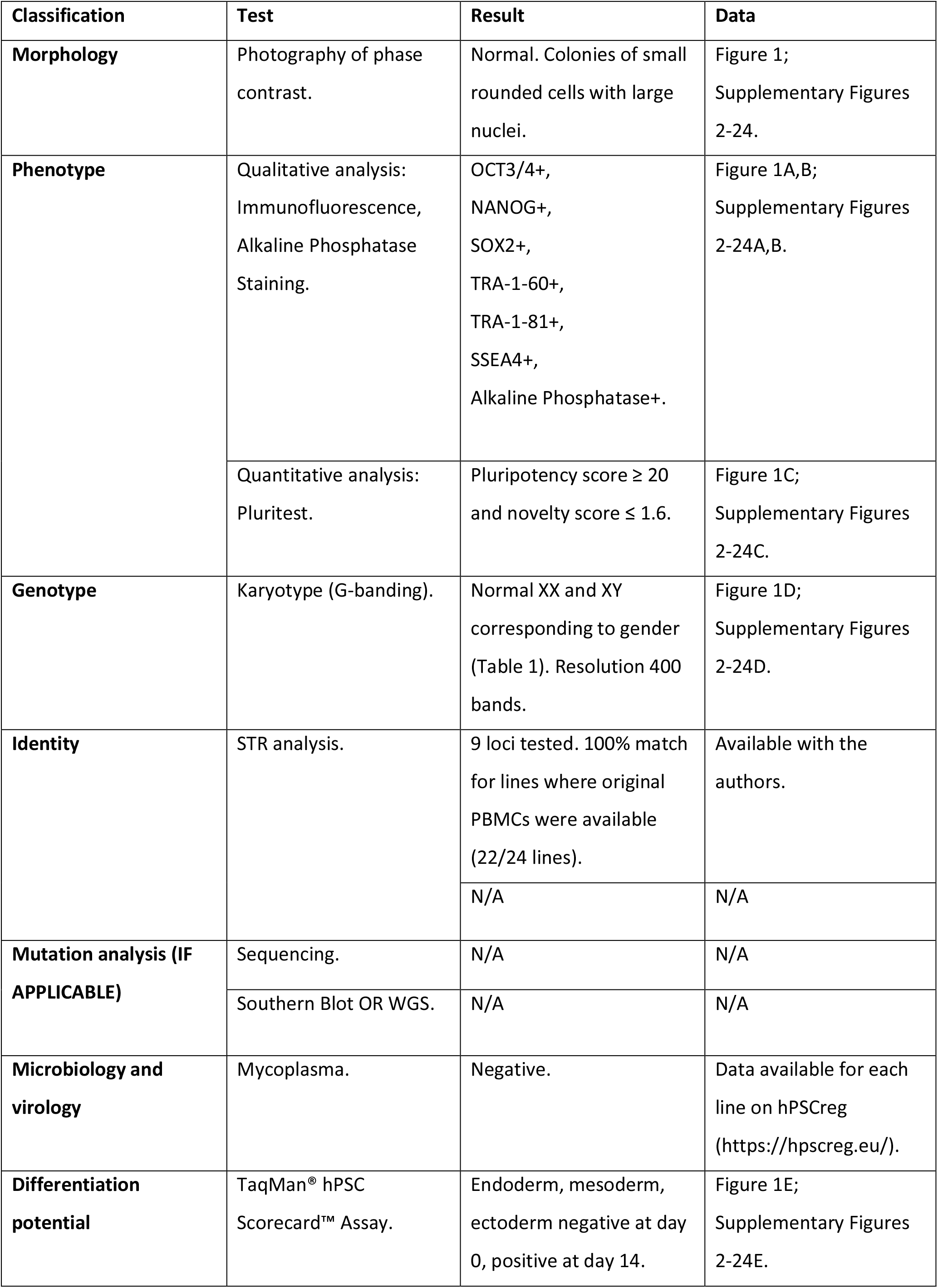
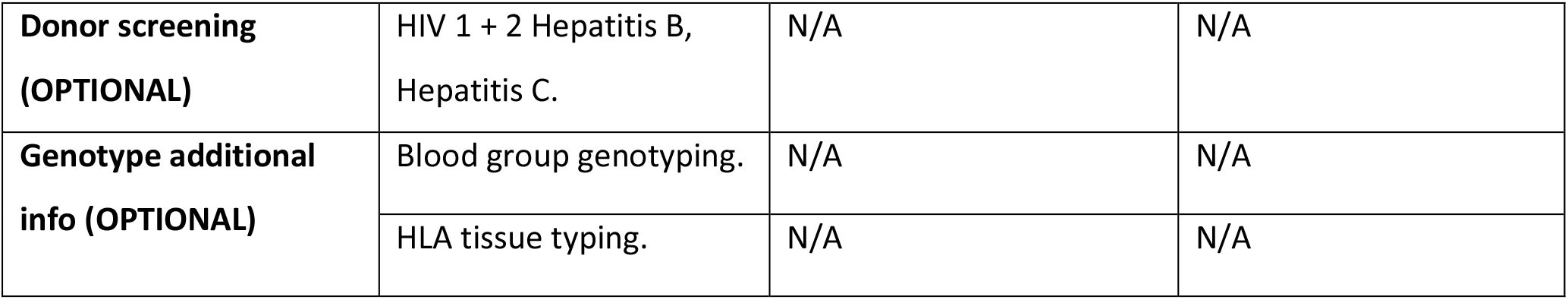
Characterization and validation

## Materials and Methods

### PBMC isolation

Blood samples were collected with NHS Lothian Research Ethics Committee Approval (10/S1103/10). Blood samples were collected in Sodium Citrate BD Vacutainer CPT tubes (BD, Cat. 362761) (three tubes per participant). For samples EDi021-A, EDi025-A, EDi028-A, EDi030-A, EDi031-A, EDi032-A, EDi033-A, EDi034-A, and EDi035-A PBMC isolation was performed by Roslin Cells. For all other lines, PBMC isolation was performed by the Edinburgh Clinical Research Facility (ECRF).

### Generation of human iPSCs

Generation of human iPSCs lines from PBMCs was performed using nucleofection of episomal plasmids containing POU5F1, SOX2, KLF4, LIN28, L-MYC, TP53shRNA, and SV40LT.

Briefly, ~5×10^6^ cells per nucleofection of PBMCs were nucleofected with plasmid mixture 5p on the Amaxa Nucleofector 2D Device with the Amaxa Human T-cell Nucleofector® Kit. Each transfection contained the following seven factors: OCT4, SOX2, KLF4, LMYC, LIN28, SV40LT and p53 shRNA. These were delivered on the following plasmids, together with an EBNA1 plasmid for episomal plasmid maintenance: pEP4 E02S ET2K, pCXLE-hOCT3/4-shp53-F, pCXLEhUL, pCXLE-hSK, and pCXWB-EBNA1. Nucleofected cells were immediately seeded into three wells of a 6-well plate covered with mitomycin treated mouse embryonic feeder (MEF) layer, and fed with 2mL of either αβ T-cell medium (X-vivo10 supplemented with 30U/ml IL-2 and 5ul/well Dynabeads Human T-activator CD3/CD28) or non T-cell medium (αMEM supplemented with 10% FBS, 10ng/ml IL-3, 10ng/ml IL-6, 10ng/ml G-CSF and 10ng/ml GM-CSF).

Two days after nucleofection, an equal amount of Primate ESC medium (ReproCell) containing 5ng/ml bFGF (for MEF condition) was added to the wells without aspirating the previous medium. Beginning on day four, the medium was gently aspirated from each well and 2mL of the appropriate fresh reprogramming media was added to each well. Medium was replaced every other day. At approximately day 18 post nucleofection, individual colonies were observed in all wells of each condition. Individual PBMC-iPSC colonies with ES/iPSC-like morphology appeared between day 25-32 and those with best morphology were mechanically isolated, transferred onto 12-well plates with fresh Matrigel™ Matrix (Corning/BD Biosciences, Cat. 354230), and maintained in mTeSR®1 medium (STEMCELL). The iPSC clones were further expanded and scaled up for further analysis. All cultures were maintained at 20% O_2_, 5% CO_2_ during the reprogramming process.

### iPSC maintenance and storage

Human iPSCs were cultured in mTeSR®1 medium (STEMCELL) on growth factor-reduced Matrigel™ Matrix (BD Biosciences)-coated plates at 37°C in a 5% CO^2^ incubator. Briefly, 70–90% confluent human iPSC colonies were passaged chemically (Versene, Life Technologies, Cat. 15040-066 or ReLeSR, StemCell Technologies, Cat. 05872) or mechanically by StemPro® EZPassage™ Disposable Stem Cell Passaging Tool (Life Technologies, Cat. 23181-010) and re-plated at a 1:6 or 1:9 ratio depending on the cell line. The iPSCs were expanded for 6-22 passages during which period various characterization assays were performed. The iPSCs were cryopreserved using CryoStor CS10 (StemCell Technologies) and an isopropanol freezing vessel at −80°C overnight. The cryopreserved vials were subsequently stored in liquid nitrogen tanks for long-term storage. Working Cell Banks (WCB) of iPSCs were cryopreserved at passage 9-14 and then Distribution Cells Banks (DCB) were created between passages 18-22.

### Mycoplasma testing

The absence of mycoplasma contamination in the iPSC lines were confirmed monthly using the MycoAlert Detection Kit, a selective biochemical test (LONZA, Cat. LT07-1188).

### EBNA-related gene analysis

Genomic DNA (250ng) was harvested from all cell lines and an embryonic stem cell line (H9) was used a negative control. Primers that recognize EBNA1 along with housekeeping gene Glyceraldehyde 3-phosphate dehydrogenase (GAPDH), which was used as a housekeeping gene, were included in this study (Table 2). PCR was run for 35 cycles at 95°C for 30 seconds, 60°C for 30 seconds, and 72°C for 30 seconds.

### TCRB and TCRG T-Cell Clonality Assay

TCRB and TCRG T-Cell Clonality testing was conducted using Gene Rearrangement and Translocation assays from Invivoscribe Technologies, Inc. Genomic DNA was harvested from all iPSC lines using the MagMAX^TM^ DNA Multi-Sample Ultra 2.0 Kit (Cat. A36570) from Applied Biosystems and it was re-suspended to a final concentration of 100-400μg per ml in dilution buffer. Three Clonal Control DNA and one Polyclonal Control DNA provided with the kit were used. PCR was carried out as per the manufacturer’s protocol. PCR products were analysed using 6% TBE gel electrophoresis with gel red staining.

### Karyotyping

Human PBMC-iPSCs were incubated in Colcemid (100ng/mL; Life Technologies) for 30 minutes at 37°C and then dissociated using TrypLE for 5 minutes. They were then washed in phosphate buffered saline (PBS) and incubated at 37°C in 5mL hypotonic solution (1g KCl, 1g Na Citrate in 400mL water) for 30 minutes. The cells were centrifuged for 2.5 minutes at 1500RPM and re-suspended in fixative (methanol: acetic acid, 3:1) at room temperature for 5 minutes. This was repeated twice, and finally cells were re-suspended in 500μl of fixative solution and submitted to the Cedars-Sinai Clinical Cytogenetics Core for G-band karyotyping. Karyotyping of each iPSC line was conducted at early and late passage, between passages 6-22.

### Immunocytochemistry

iPSCs were plated on Matrigel™ Matrix (BD Biosciences)-coated glass coverslips or optical-bottom 96-well plates (Thermo, Cat. 165305) and subsequently fixed in 4% paraformaldehyde (ten minutes, room temperature (RT)). All cells were blocked in 4-5% goat or donkey serum with 0.1% Triton X-100 (one hour, RT) and incubated with primary antibodies (Table 3) for either 3 hours at RT or overnight at 4°C. Cells were then rinsed and incubated in species-specific AF488 or AF594-conjugated secondary antibodies (1:500) (one hour, RT), followed by DAPI (0.5-1μg/ml; Sigma) to counterstain nuclei (10 minutes, RT). Cells were imaged using Nikon/Leica microscopes or Image Express. The iPSCs exhibited an embryonic stem cell like morphology, and expressed a range of pluripotency markers (OCT3/4, NANOG, SOX2, TRA-1-60, TRA-1-81, SSEA4) (Figure 1B, Supplementary Figures 2-24B).

**Table 3:**
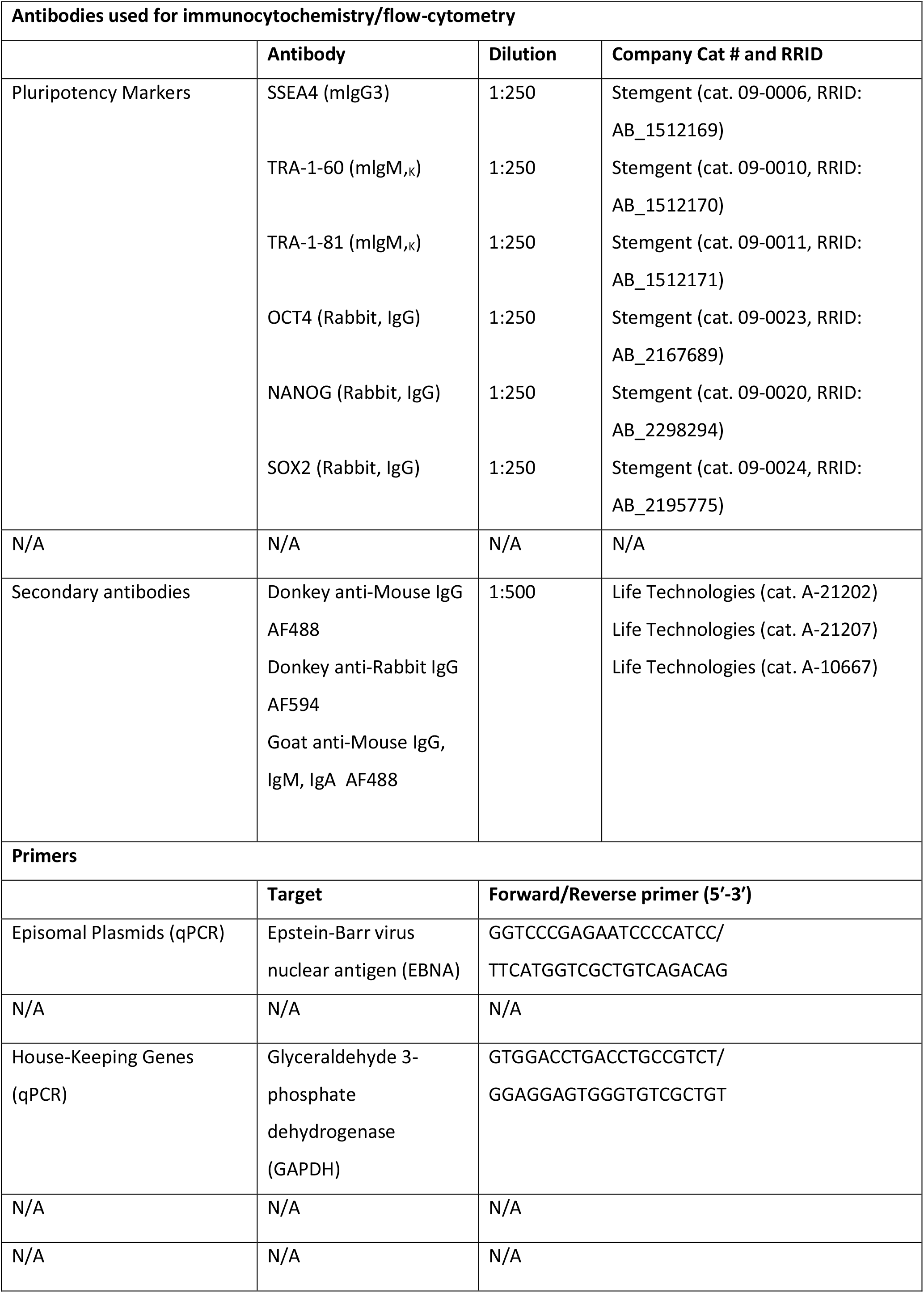
Reagents details

### Alkaline phosphatase staining

Alkaline phosphatase staining was performed using the Alkaline Phosphatase Staining Kit II (Stemgent, Cat. 00-0055) according to the manufacturer’s instructions.

### PluriTest

PluriTest was used to assess the pluripotency of undifferentiated iPSCs (Figure 1C, Supplementary Figures 2-24C). Cell pellets were sent to Life Technologies Corporation for the PluriTest Service. Total RNA was isolated using the PureLinkTM RNA Mini Kit (Thermo Fisher Scientific) and quantified using NanoDropTM. 100ng total RNA was used to prepare the GeneChip® for the PluriTest™. In this assay, 36,000 transcripts and variants against a >450 sample reference set are assessed for gene expression analysis. A non-iPSC sample was used in this experiment to serve as a control for non-pluripotency. The transcriptome of all samples were analysed and processed in the PluriTest™ algorithm to generate a pluripotency and novelty score. These two scores determine the pluripotency signature of the cell line which is represented in the pluripotency plot. The threshold for pluripotency was >20, and the threshold for novelty was <1.6.

### hPSC Scorecard Data Analysis

Applied Biosystems TaqMan®hPSC Scorecard^TM^ Panel (Thermo Fisher Scientific) was used as an additional technique to assess pluripotency and tri-lineage differentiation potential of iPSC lines using real-time qPCR assays (Figure 1E, Supplementary Figures 2-24E). Total RNA from undifferentiated and EB differentiated iPSC lines was isolated using MagMAX^TM^ mirVana^TM^ Total RNA Isolation Kit (A27828), and 1μg of RNA was used to make cDNA using the High Capacity cDNA Reverse Transcription Kit (4368813), both from Applied Biosystems. TaqMan qRT-PCR was carried out using the hPSC Scorecard 384w Fast plate and following manufacturer protocol. We analysed the gene expression data from the TaqMan®hPSC Scorecard^TM^ Panel using the web-based hPSC Scorecard^TM^ Analysis Software (Thermo Fisher Scientific).

### Embryoid Body (EB) Formation

IPSC lines were allowed to differentiate by EB formation. Briefly, iPSCs were lifted from 3 wells of a 6 well plate using a cell scraper and seeded in a T25 flask treated with poly-HEMA to prevent cell attachment in EB media containing: IMDM basal media (Cat. 12440061), 17% KnockOut Serum Replacement (KOSR; Cat. 10828028), 1% non-essential amino acids (Cat. 11140050), 1% Antibiotic-Antimycotic (Cat. 15240062) and 110µM β-Mercaptoethanol (Cat. 21985023), all from Thermo Fisher. EBs were allowed to form by self-aggregation, grow and differentiate for 14 days in EB culture media replacing it twice a week. Differentiation to endoderm, mesoderm and ectoderm was assessed by TaqMan® hPSC Scorecard™ Assay (Figure 1E, Supplementary Figures 2-24E).

#### STR Analysis

Short Tandem Repeat (STR) Analysis is conducted to confirm iPSC genetic identity. For that, a frozen vial of the parent PBMCs and a frozen vial of the reprogramed iPSC line at late passage (18-21, depending on the cell line) are sent to IDEXX BioResearch. STR profile and interspecies contamination testing is analysed. iPSC line human authentication was conducted at IDEXX BioResearch by Cell Check^TM^. Profiling included using a nine marker STR profile (AMEL, CSF1PO, D13S317, D16S539, D5S818, D7S820, TH01, TPOX and vWA) and interspecies contamination check for human, mouse, rat, African green monkey and Chinese hamster cells. Comparative analysis was conducted between parent PBMCs and reprogrammed iPSC lines.

## Acknowledgements

We thank the LBC1936 participants who took part in this study, and gratefully acknowledge the contribution of the late Professor John M. Starr, who was the Lothian Birth Cohort 1936’s research medical doctor from its beginning in 2004 until 2018. Additionally, we would like to thank the Edinburgh Clinical Research Facility nursing staff, Roslin Cells, and the LBC research team members for their contributions to sample collection, sample processing, and data processing respectively. Finally, we thank The David and Janet Polak Foundation for their support of the Cedars-Sinai iPSC Core laboratory.

