## Supplementary Figures for "Generation of twenty four induced pluripotent stem cell lines from twenty four members of the Lothian Birth Cohort 1936"

#### Selection of iPSC line donors based on cognitive ageing

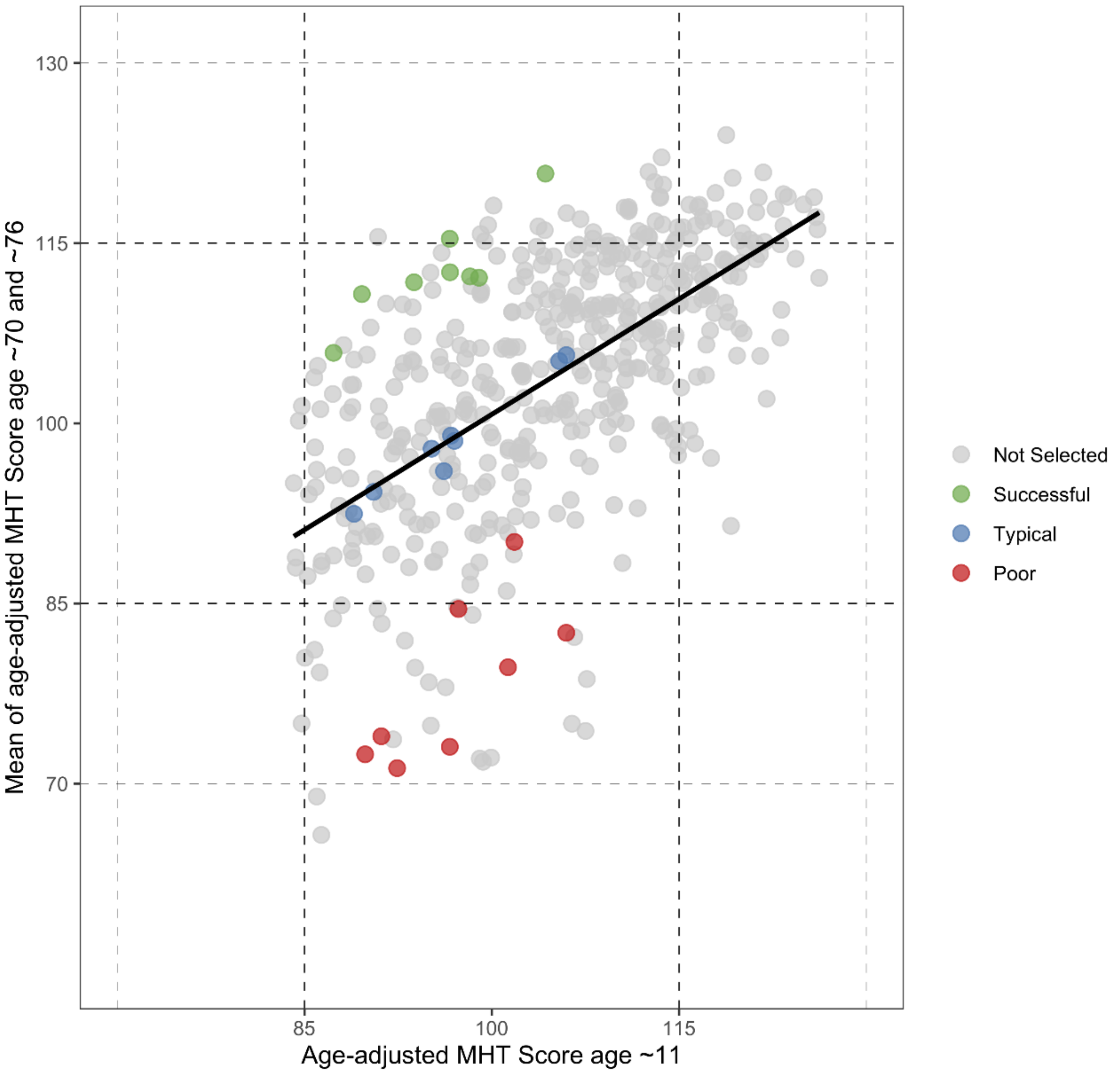


**sFig1:** Selection of iPSC candidates based on cognitive ageing profiles in the Lothian Birth Cohort 1936. Dashed lines are added to show ±1SD from the mean for the age 11 Moray House Test (MHT) score and ±1 and 2SDs from the mean for later-life MHT score.

### sFig2. Characterization for iPSC line EDi022-A

## A. AP

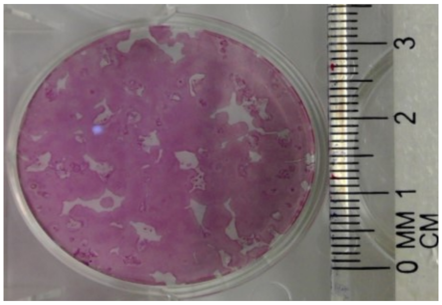


#### B. Immunocytochemistry

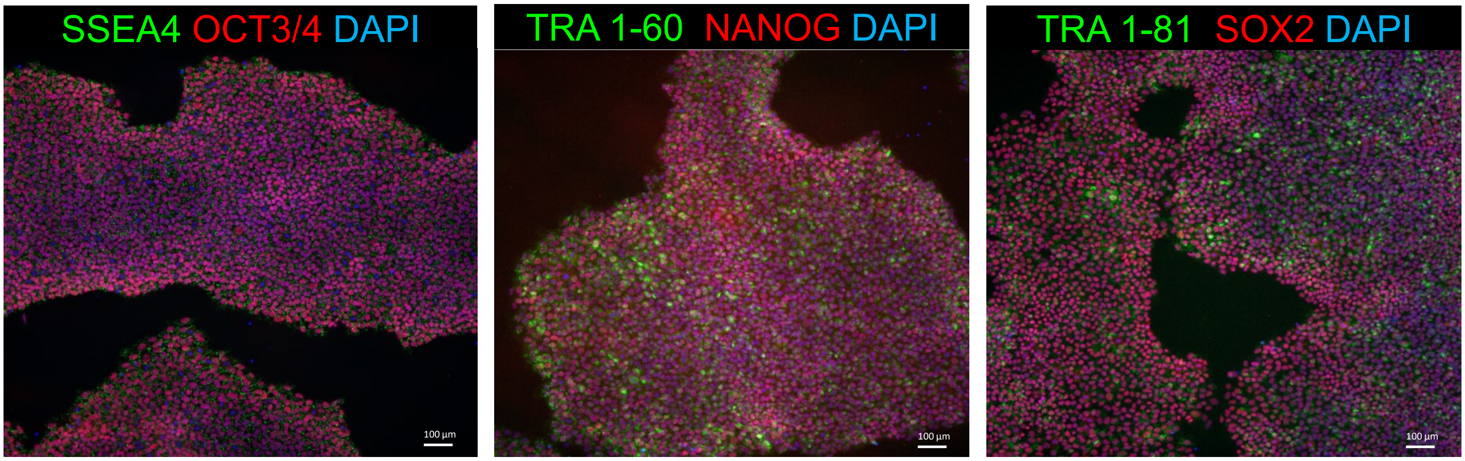


#### C. PluriTest

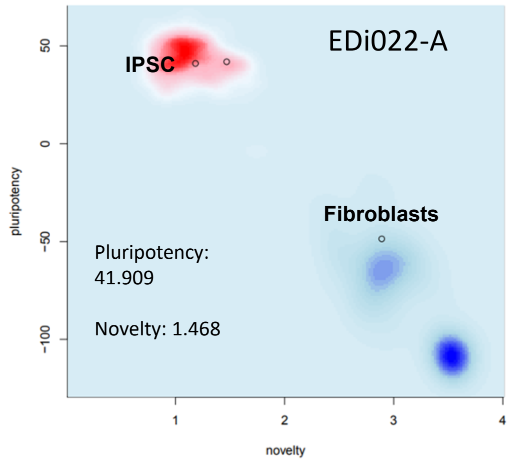


#### D. G-Band karyotype

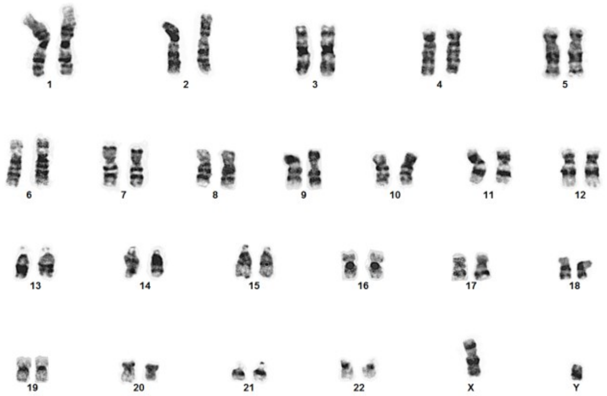


#### E. hPSC Scorecard

| iPSCs | | | | Embryoid bodies | | | |
|---|---|---|---|---|---|---|---|
| Self-renew | Ecto | Meso | Endo | Self-renew | Ecto | Meso | Endo |
| 0.29 | -0.10 | 0.62 | -0.53 | -3.06 | 1.41 | 0.76 | -0.32 |

### sFig3: Characterization for iPSC line EDi023-A

## A. AP

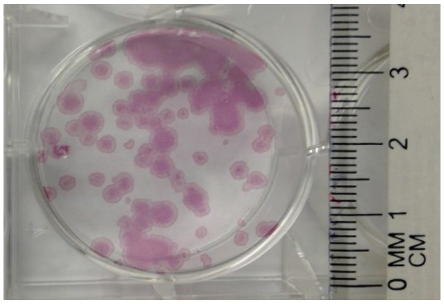


#### B. Immunocytochemistry

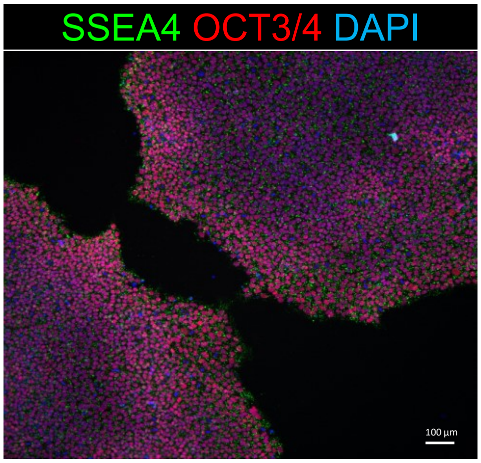


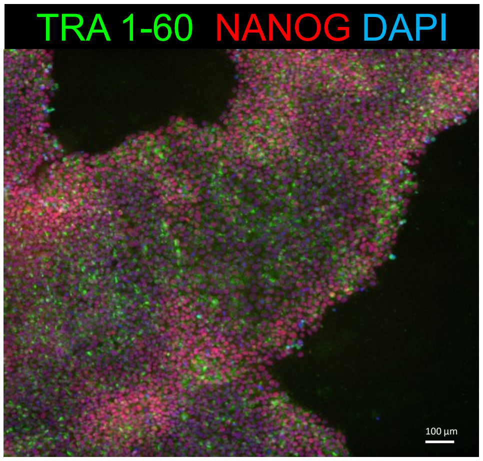


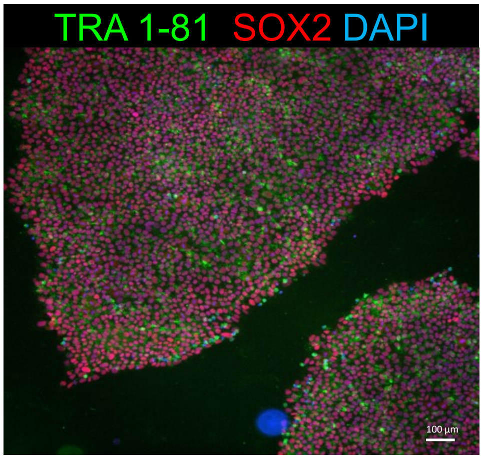


#### C. PluriTest

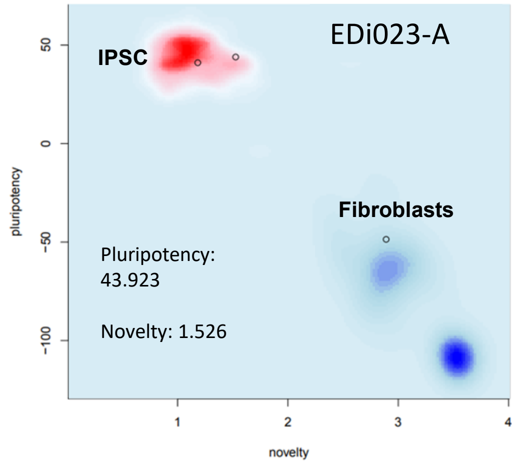


#### D. G-Band karyotype

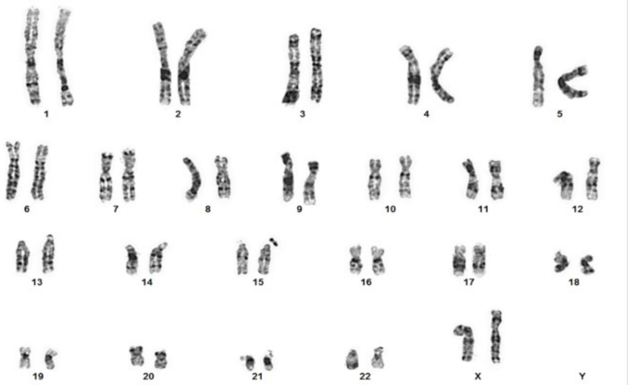


#### E. hPSC Scorecard

| iPSCs | | | | Embryoid bodies | | | |
|---|---|---|---|---|---|---|---|
| Self-renew | Ecto | Meso | Endo | Self-renew | Ecto | Meso | Endo |
| + | − | − | − | − | + | + | + |
| 0.28 | -0.30 | 0.34 | -0.67 | -3.49 | 2.38 | 3.38 | 1.12 |

### sFig4: Characterization for iPSC line EDi025-A

## A. AP

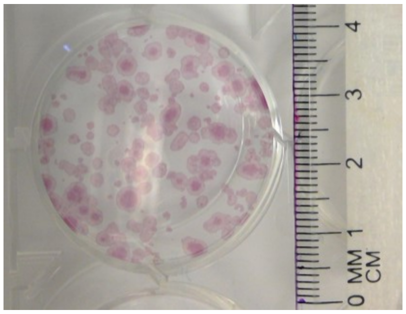


#### B. Immunocytochemistry

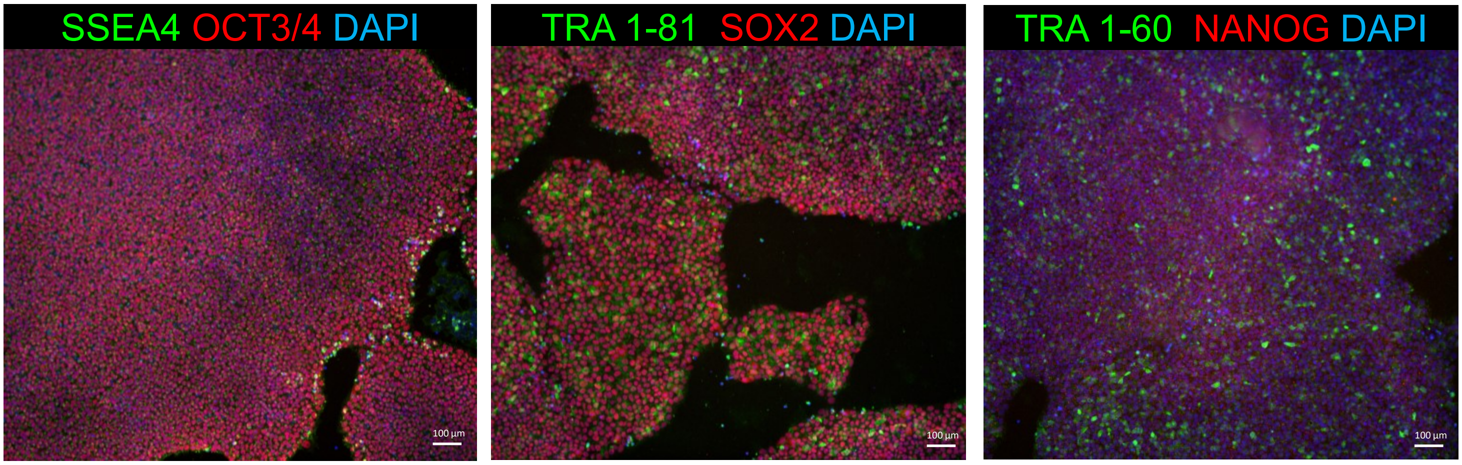


#### C. PluriTest

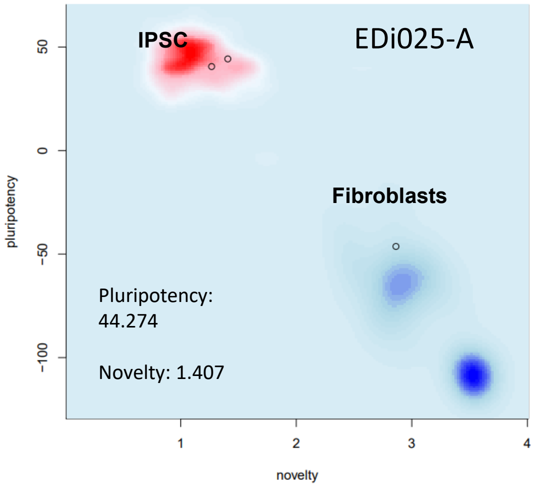


#### D. G-Band karyotype

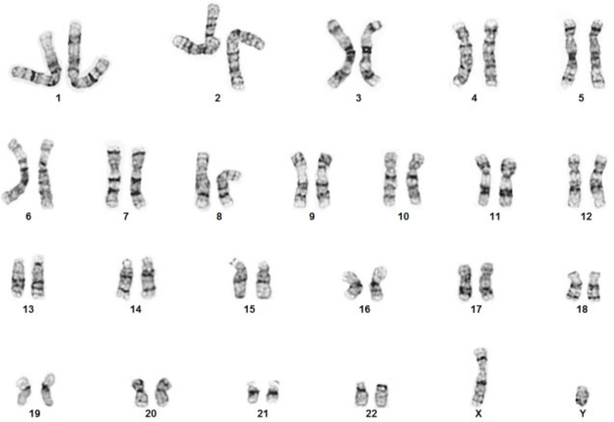


#### E. hPSC Scorecard

| iPSCs | | | | Embryoid Bodies | | | |
|---|---|---|---|---|---|---|---|
| Self-renew | Ecto | Meso | Endo | Self-renew | Ecto | Meso | Endo |
| + | − | − | − | − | + | + | + |
| -0.01 | -0.33 | -0.39 | -1.15 | -7.29 | 1.49 | 2.19 | 0.57 |

### sFig5: Characterization for iPSC line EDi026-A

## A. AP

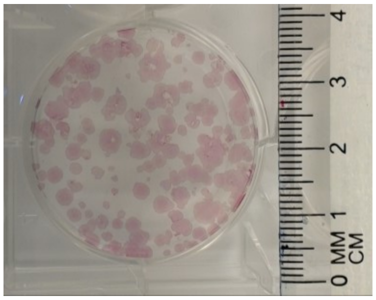


#### B. Immunocytochemistry

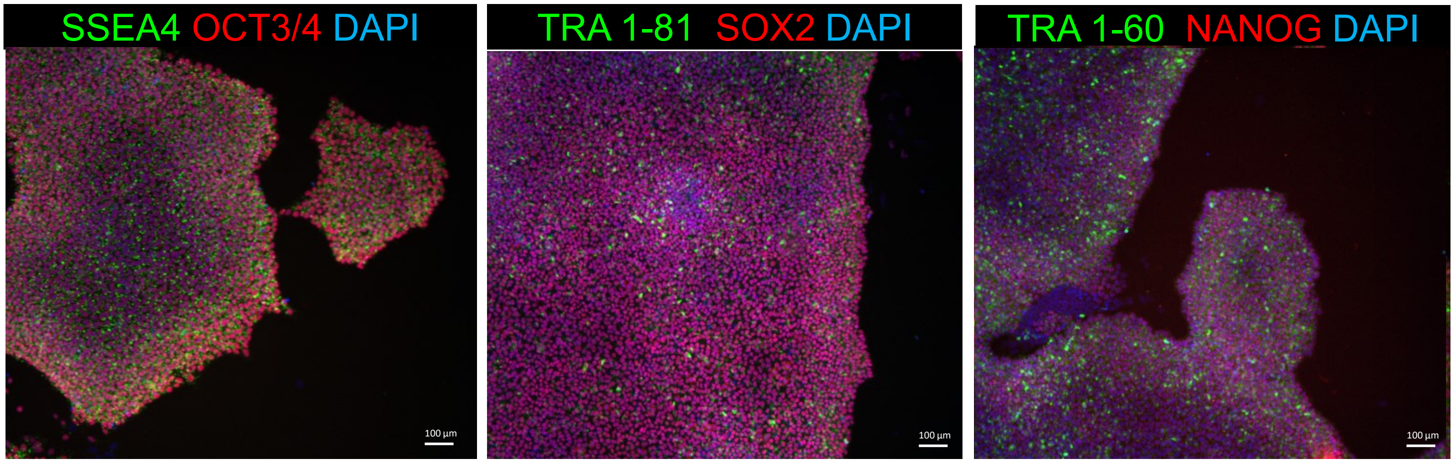


#### C. Pluritest

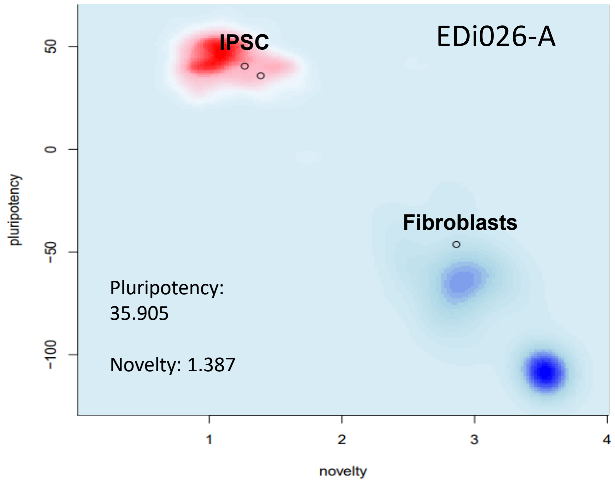


#### D. G-Band karyotype

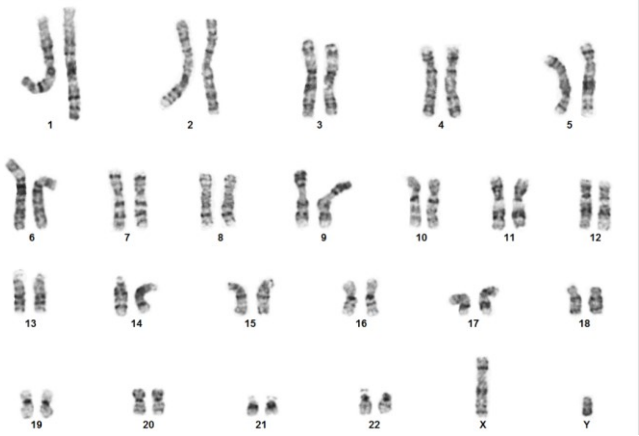


#### E. hPSC Scorecard

| iPSCs | | | | Embryoid Bodies | | | |
|---|---|---|---|---|---|---|---|
| Self-renew | Ecto | Meso | Endo | Self-renew | Ecto | Meso | Endo |
| + | − | − | − | − | + | + | + |
| -0.41 | 0.50 | 0.42 | -0.83 | -6.72 | 1.74 | 1.19 | 0.69 |

### sFig6: Characterization for iPSC line EDi027-A

## A. AP

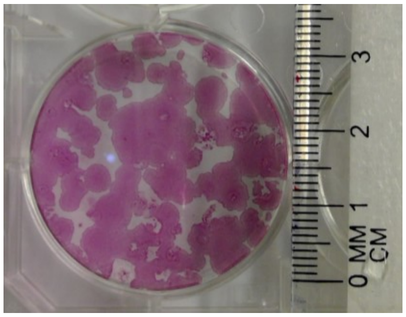


#### B. Immunocytochemistry

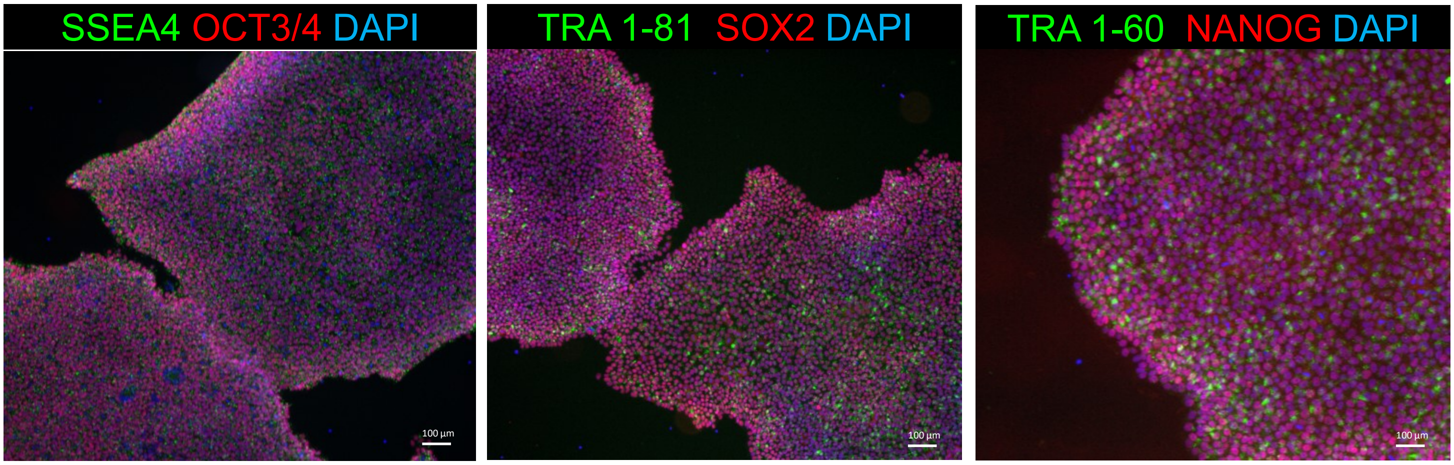


#### C. Pluritest

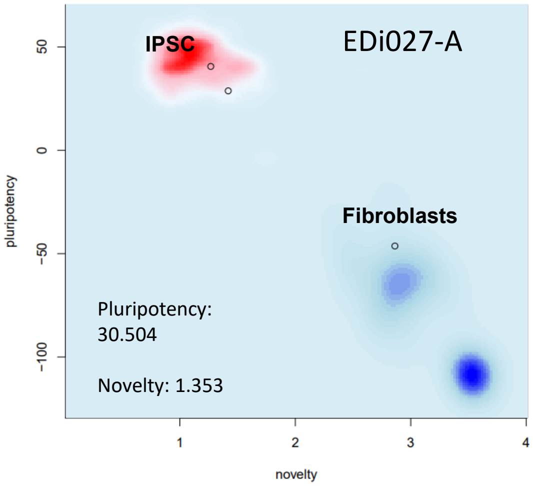


#### D. G-Band karyotype

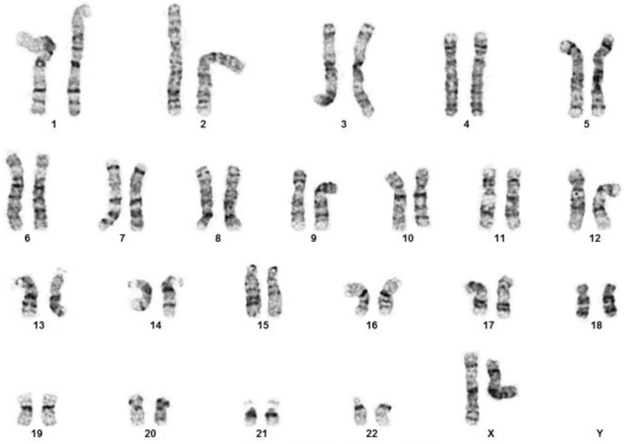


#### E. hPSC Scorecard

| IPSCs | | | | Embryoid Bodies | | | |
|---|---|---|---|---|---|---|---|
| Self-renew | Ecto | Meso | Endo | Self-renew | Ecto | Meso | Endo |
| + | O | − | − | − | + | + | + |
| 0.18 | 0.61 | 0.10 | -0.75 | -6.07 | -1.72 | -3.29 | -1.78 |

### sFig7: Characterization for iPSC line EDi028-A

## A. AP

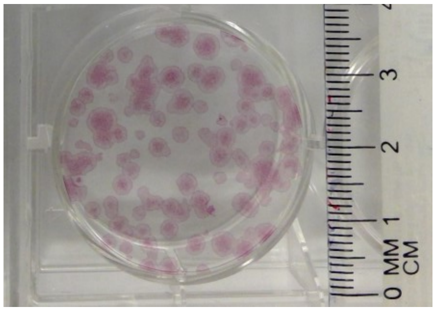


#### B. Immunocytochemistry

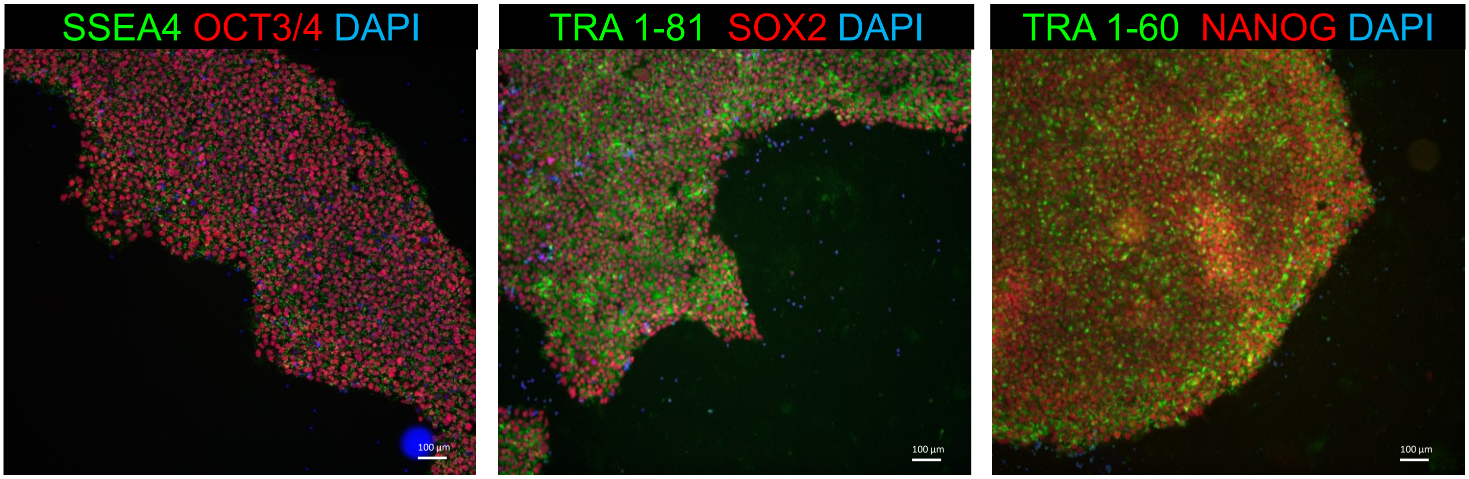


#### C. Pluritest

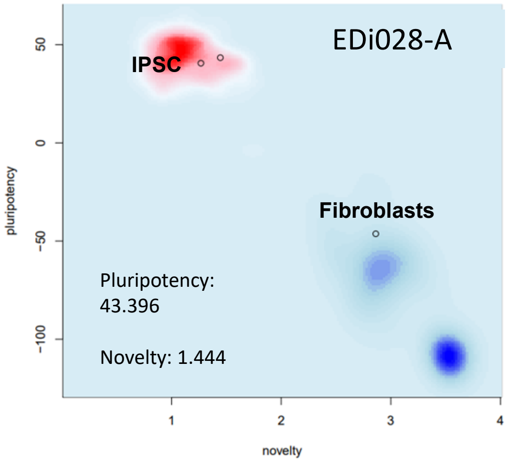


#### D. G -Band karyotype

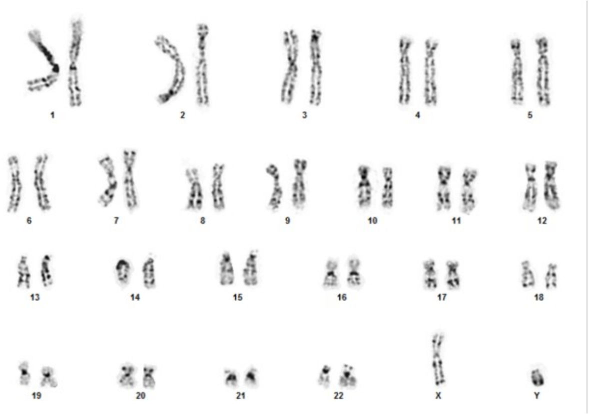


#### E. hPSC Scorecard

| IPSCs | | | | Embryoid Bodies | | | |
|---|---|---|---|---|---|---|---|
| Self-renew | Ecto | Meso | Endo | Self-renew | Ecto | Meso | Endo |
| + | − | − | − | − | + | + | + |
| -0.17 | -0.17 | 0.69 | -1.00 | -6.84 | 2.14 | 2.10 | 0.69 |

### sFig8: Characterization for iPSC line EDi029-A

## A. AP

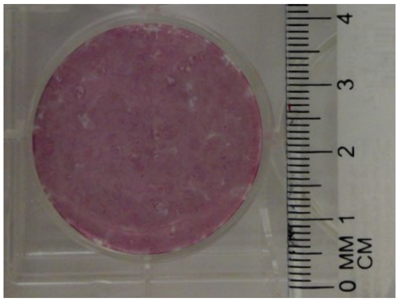


#### B. Immunocytochemistry

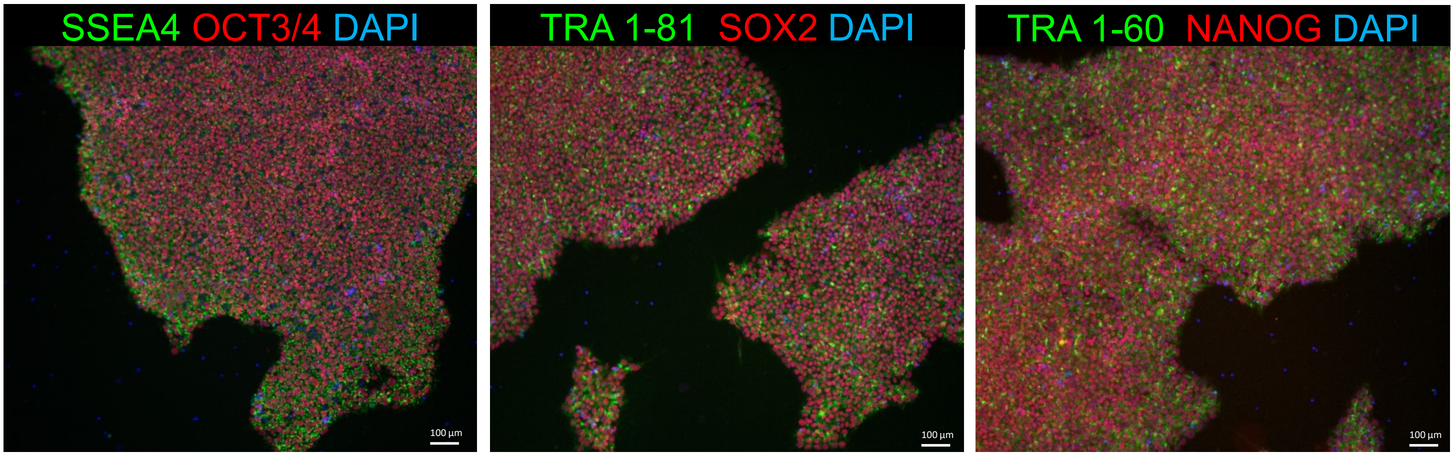


#### C. Pluritest

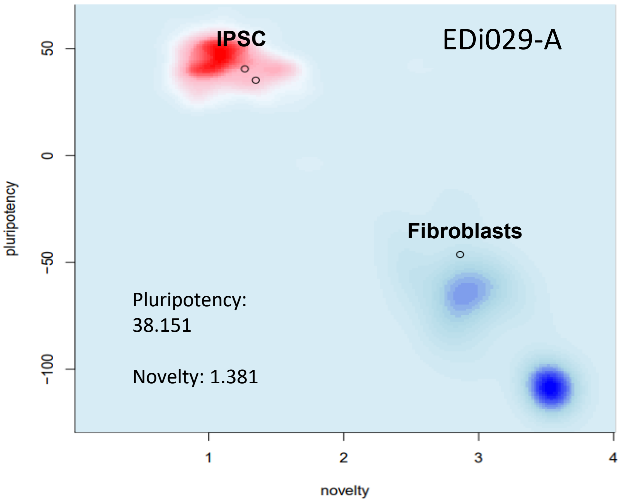


#### D. G-Band karyotype

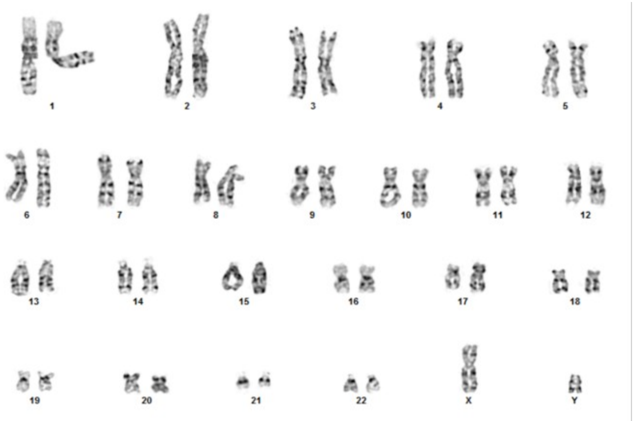


#### E. hPSC Scorecard

| iPSCs | | | | Embryoid Bodies | | | |
|---|---|---|---|---|---|---|---|
| Self-renew | Ecto | Meso | Endo | Self-renew | Ecto | Meso | Endo |
| + | − | − | − | − | + | + | + |
| 0.75 | 0.33 | 0.37 | -0.81 | -5.16 | 2.13 | 3.91 | 1.96 |

### sFig9: Characterization for iPSC line EDi030-A

## A. AP

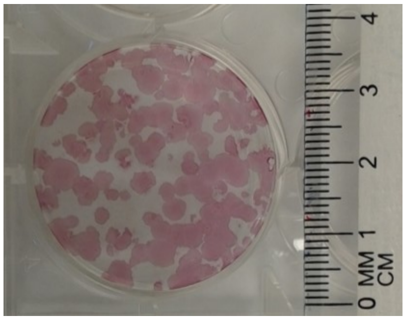


#### B. Immunocytochemistry

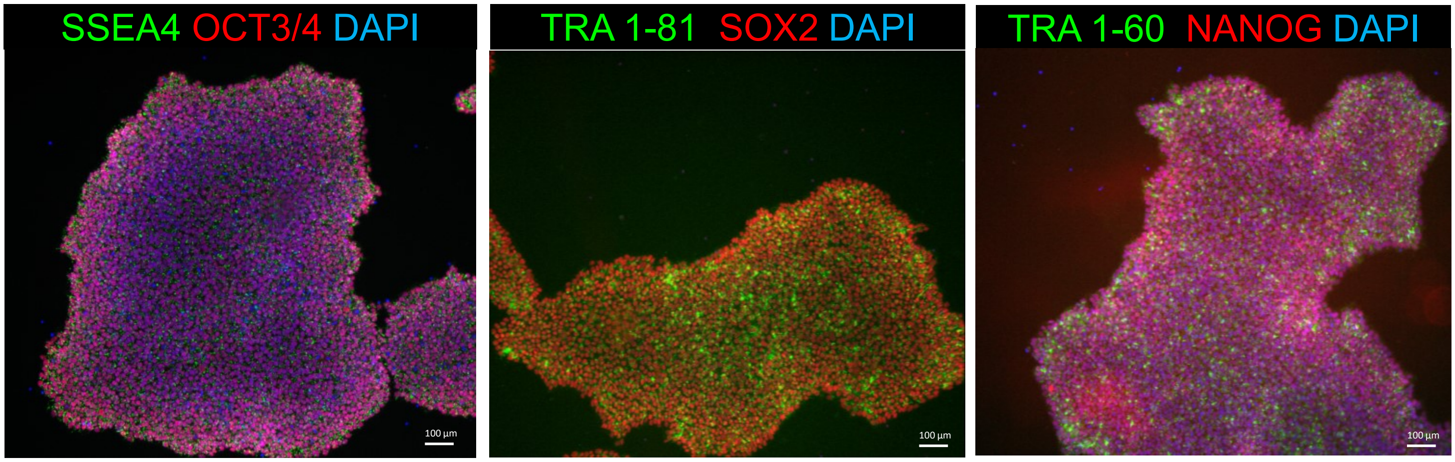


#### C. Pluritest

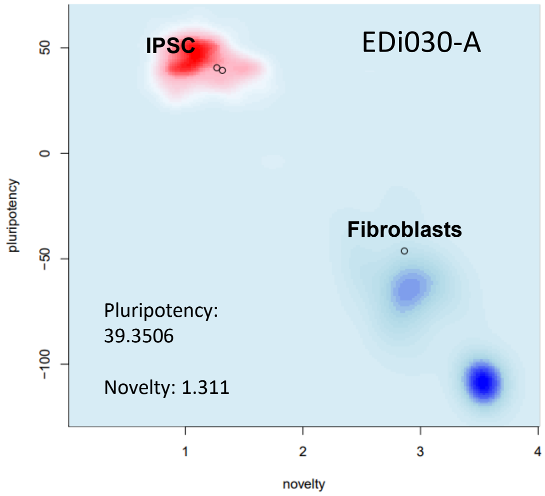


#### D. G-Band karyotype

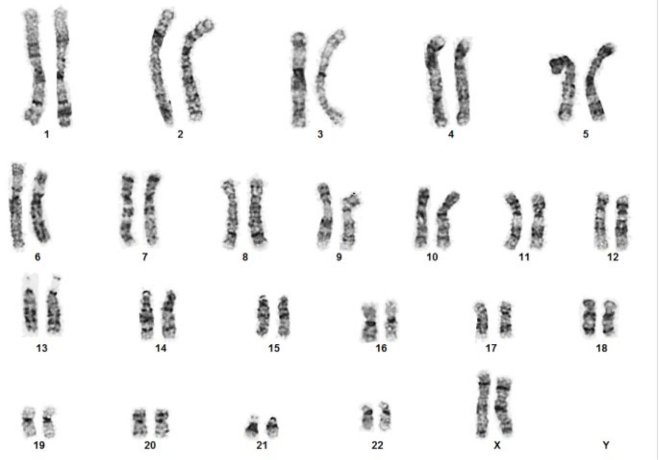


#### E. hPSC Scorecard

| IPSCs | | | | Embryoid Bodies | | | |
|---|---|---|---|---|---|---|---|
| Self-renew | Ecto | Meso | Endo | Self-renew | Ecto | Meso | Endo |
| + | − | − | − | − | + | + | + |
| -0.30 | 0.07 | 0.54 | -1.48 | -5.34 | 1.72 | 5.14 | 1.95 |

### sFig10: Characterization for iPSC line EDi031-A

## A. AP

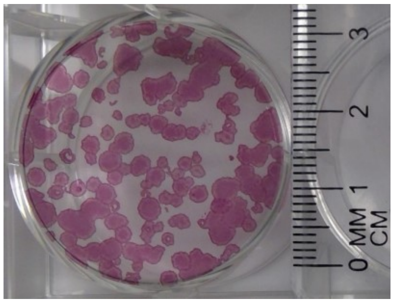


#### B. Immunocytochemistry

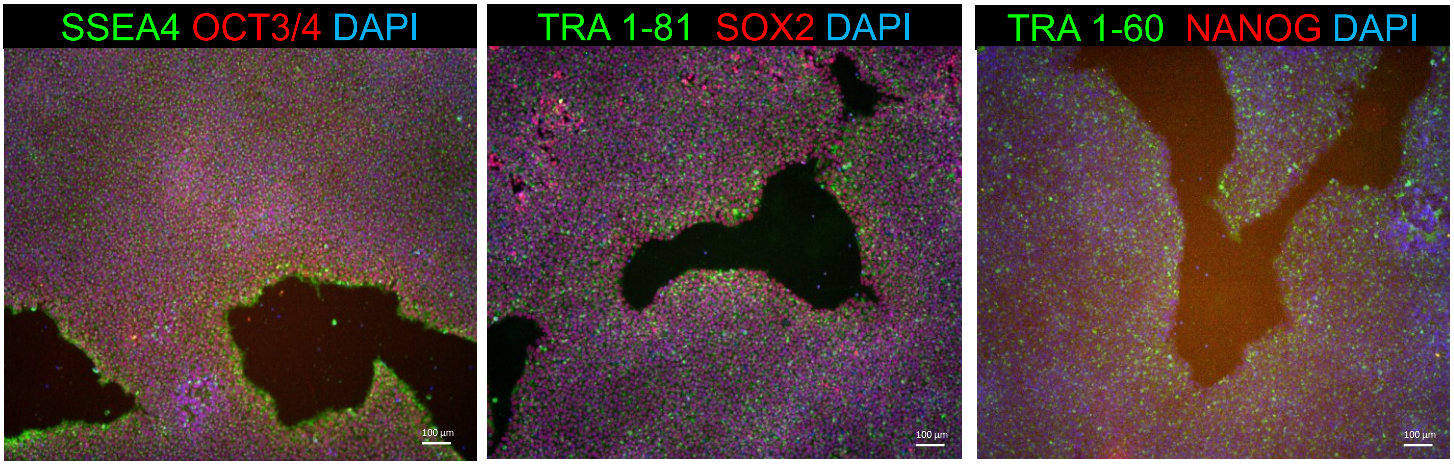


#### C. Pluritest

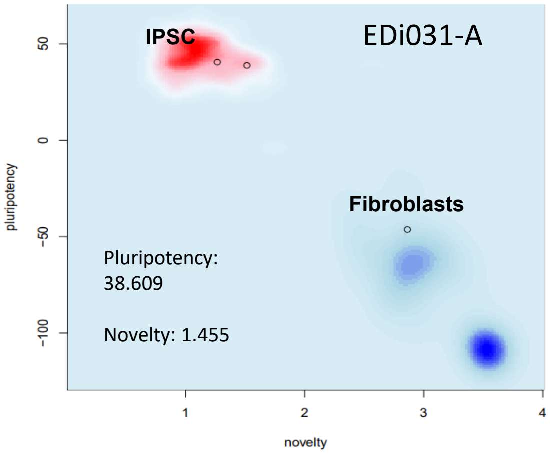


#### D. G-Band karyotype

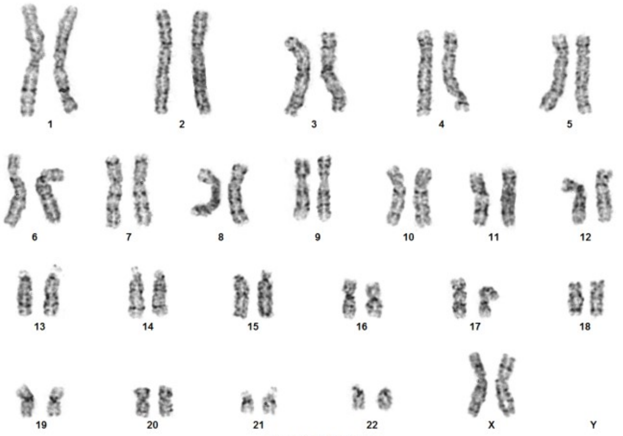


#### E. hPSC Scorecard

| iPSCs | | | | Embryoid Bodies | | | |
|---|---|---|---|---|---|---|---|
| Self-renew | Ecto | Meso | Endo | Self-renew | Ecto | Meso | Endo |
| + | − | − | − | − | + | + | + |
| -0.19 | -0.46 | -0.24 | -1.30 | -6.33 | 1.77 | 2.65 | 1.65 |

#### sFig11: Characterization for iPSC line EDi032-A

### A. AP

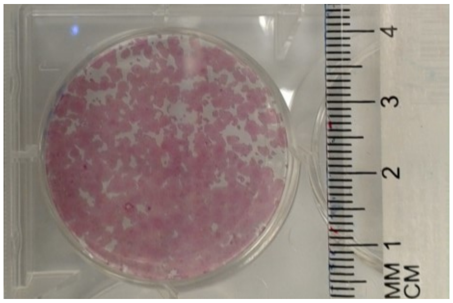


##### B. Immunocytochemistry

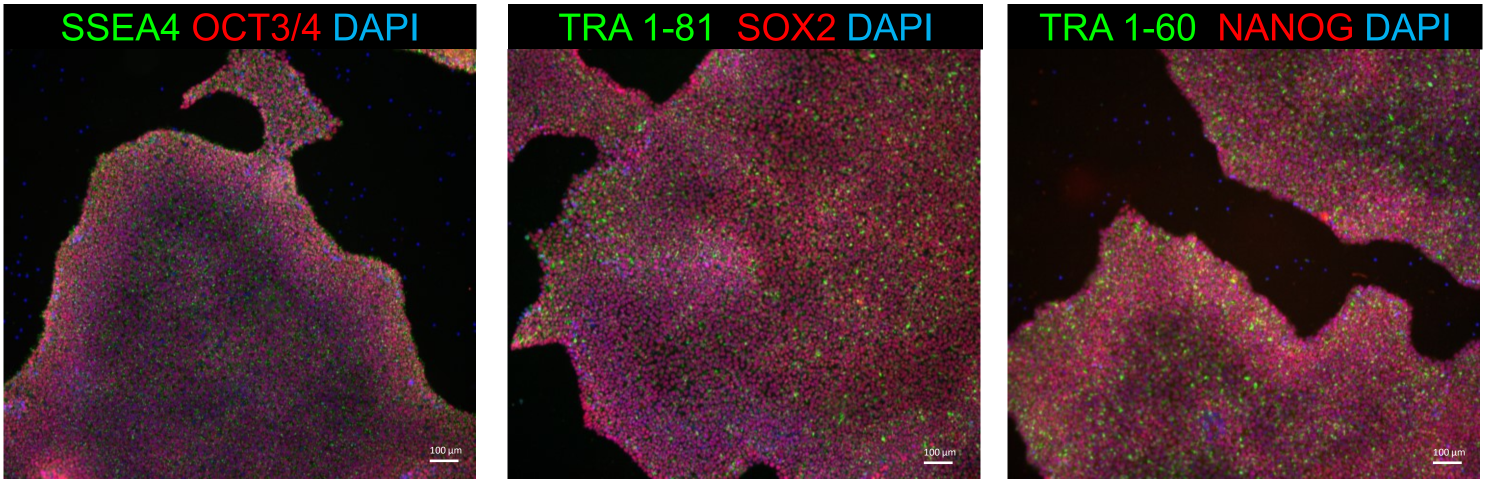


##### C. Pluritest

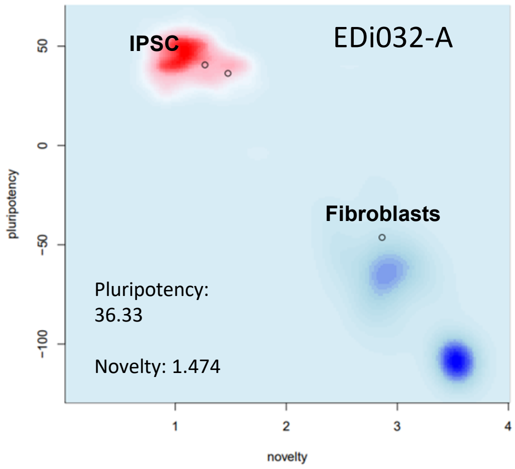


##### D. G -Band karyotype

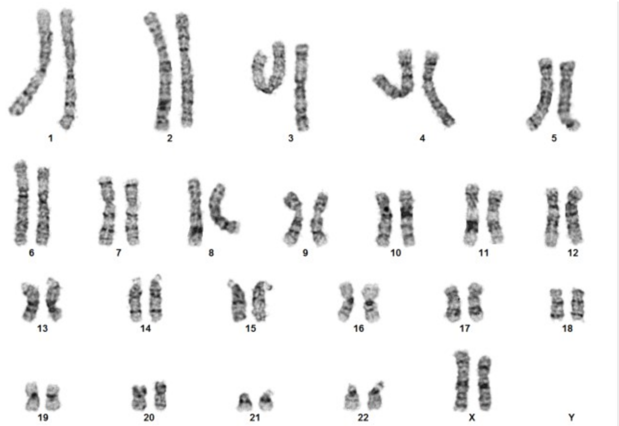


##### E. hPSC Scorecard

| IPSCs | | | | Embryoid Bodies | | | |
|---|---|---|---|---|---|---|---|
| Self-renew | Ecto | Meso | Endo | Self-renew | Ecto | Meso | Endo |
| + | − | − | − | − | + | + | + |
| 0.24 | -0.06 | 0.08 | -1.36 | -6.85 | 1.85 | 2.34 | 0.78 |

### sFig12: Characterization for iPSC line EDi033-A

## A. AP

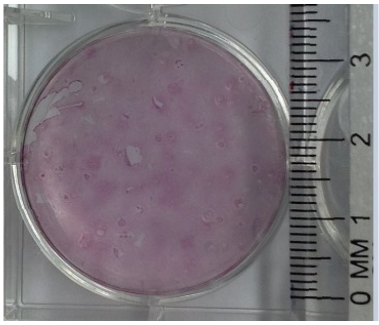


#### B. Immunocytochemistry

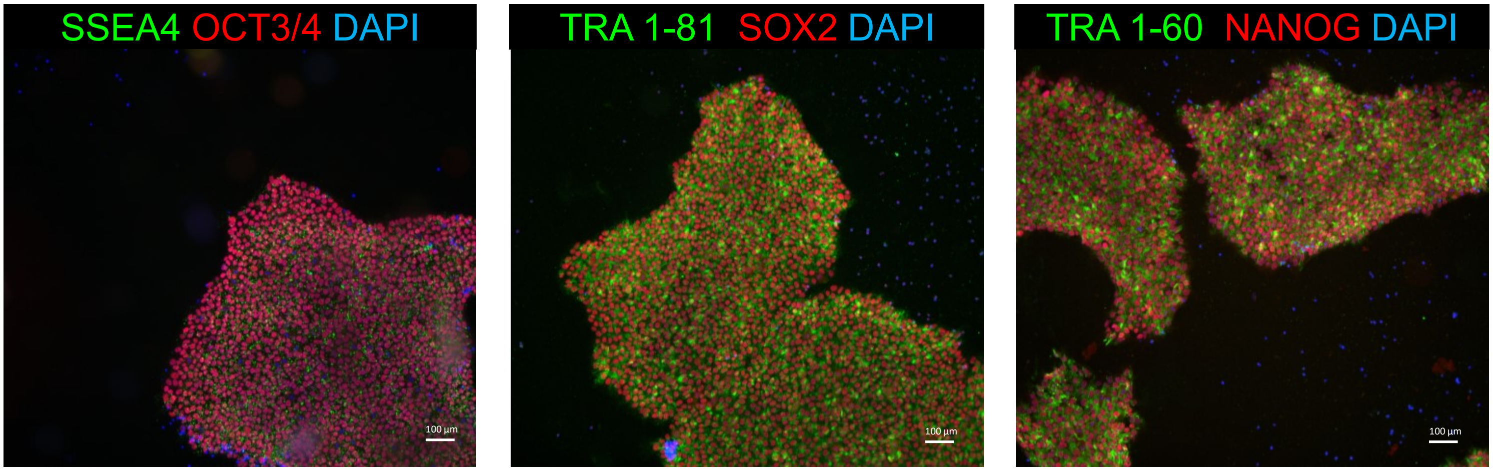


#### C. Pluritest

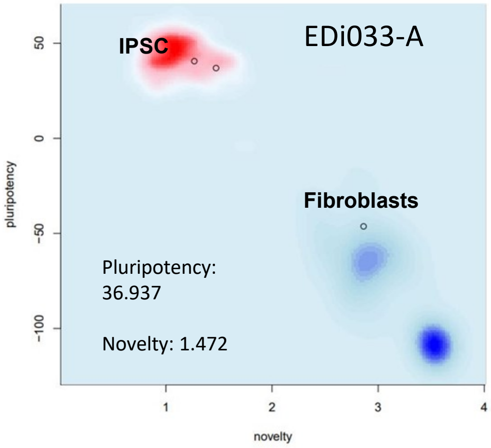


#### D. G -Band karyotype

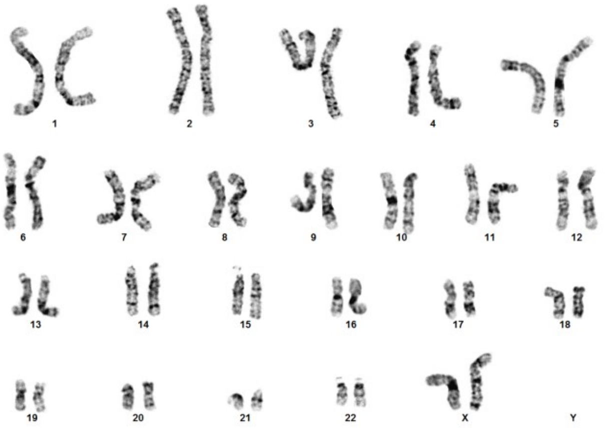

#### E. hPSC Scorecard

| IPSCs | | | | Embryoid Bodies | | | |
|---|---|---|---|---|---|---|---|
| Self-renew | Ecto | Meso | Endo | Self-renew | Ecto | Meso | Endo |
| + | − | − | − | − | + | + | + |
| 0.23 | -0.02 | -0.18 | -1.03 | -3.58 | 1.77 | 3.93 | 1.85 |

### sFig13: Characterization for iPSC line EDi034-A

## A. AP

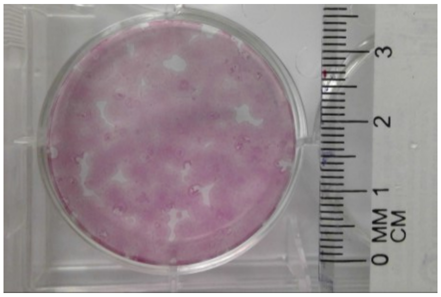


#### B. Immunocytochemistry

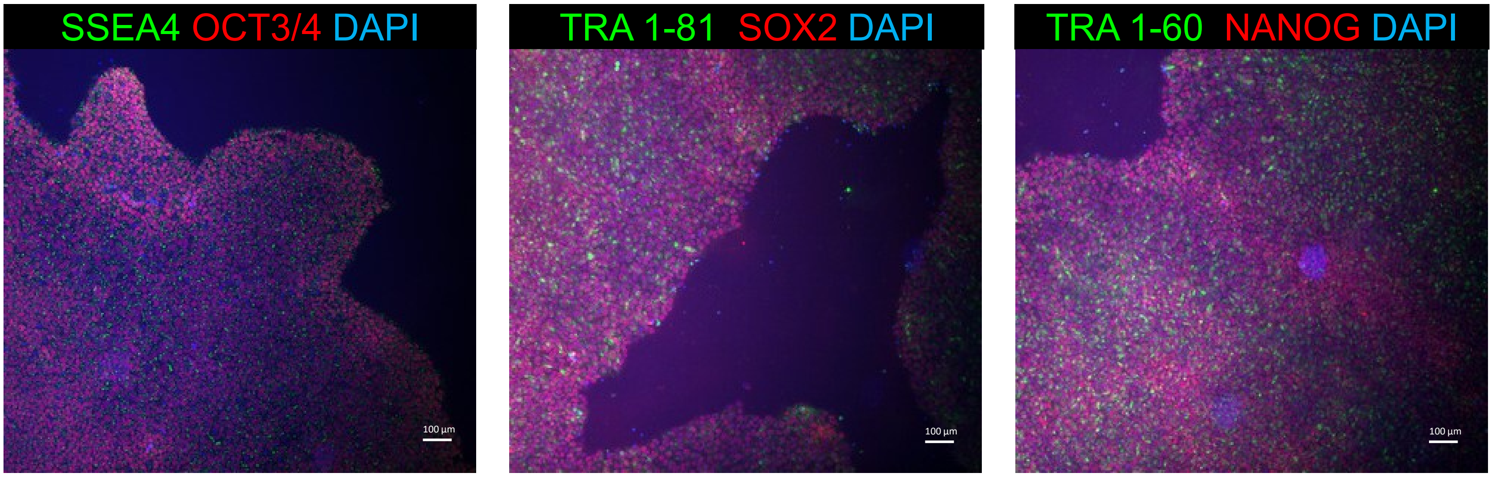


#### C. Pluritest

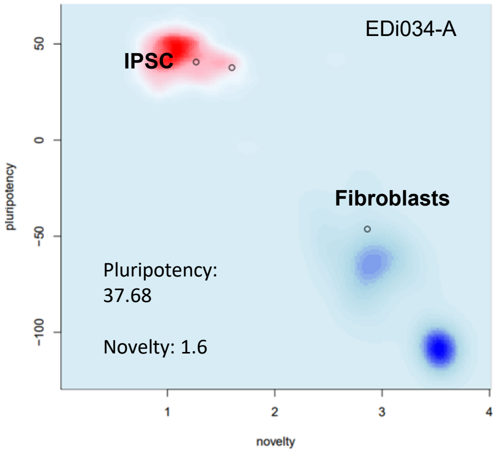


#### D. G -Band karyotype

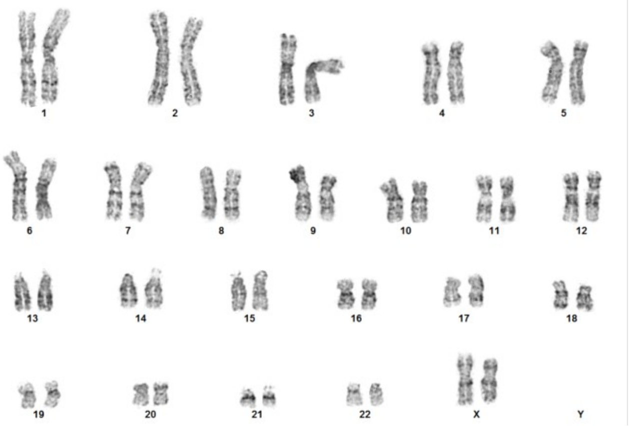


#### E. hPSC Scorecard

| IPSCs | | | | Embryoid Bodies | | | |
|---|---|---|---|---|---|---|---|
| Self-renew | Ecto | Meso | Endo | Self-renew | Ecto | Meso | Endo |
| + | − | − | − | − | + | + | + |
| 0.77 | -0.25 | -0.22 | -1.57 | -4.24 | 2.56 | 3.43 | 1.67 |

### sFig14: Characterization for iPSC line EDi035-A

## A. AP

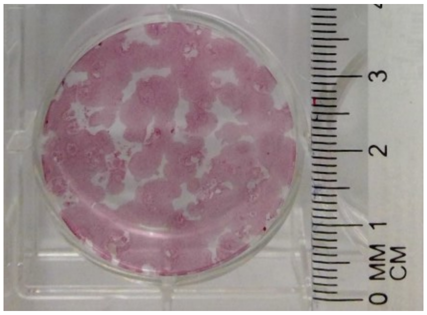


#### B. Immunocytochemistry

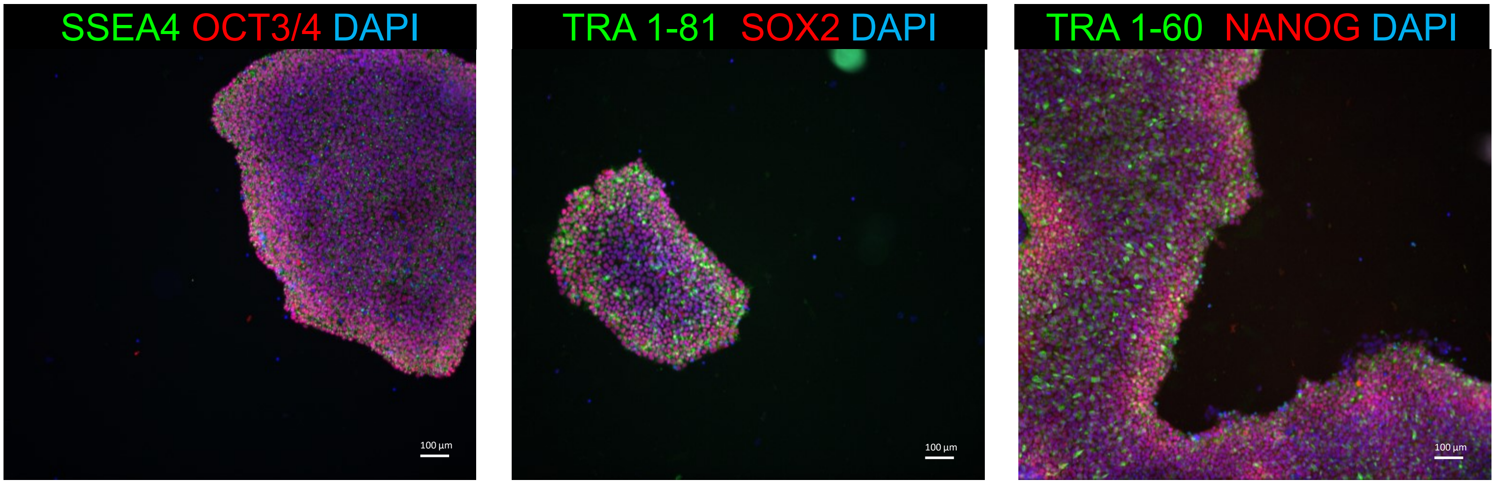


#### C. Pluritest

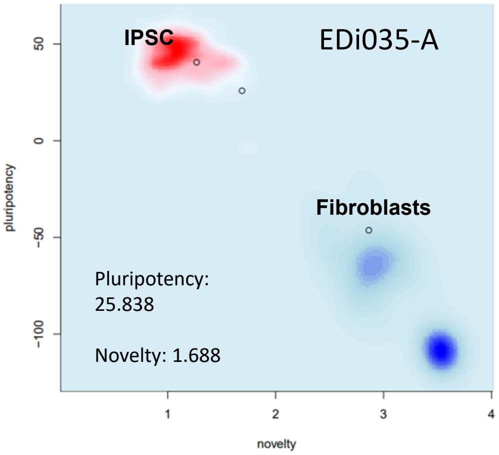


#### D. G -Band karyotype

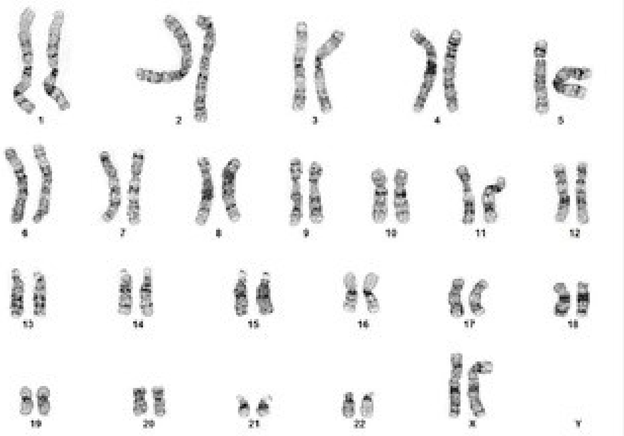

#### E. hPSC Scorecard

| iPSCs | | | | Embryoid Bodies | | | |
|---|---|---|---|---|---|---|---|
| Self-renew | Ecto | Meso | Endo | Self-renew | Ecto | Meso | Endo |
| + | − | − | − | − | + | − | O |
| 0.39 | 0.19 | 0.02 | -1.22 | -6.69 | 1.81 | 0.64 | 0.29 |

### sFig15: Characterization for iPSC line EDi036-A

## A. AP

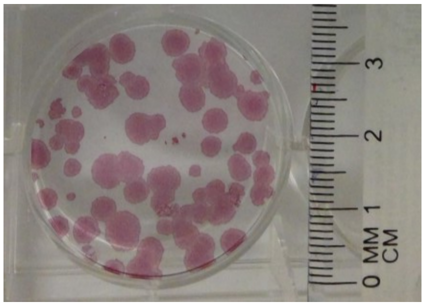


#### B. Immunocytochemistry

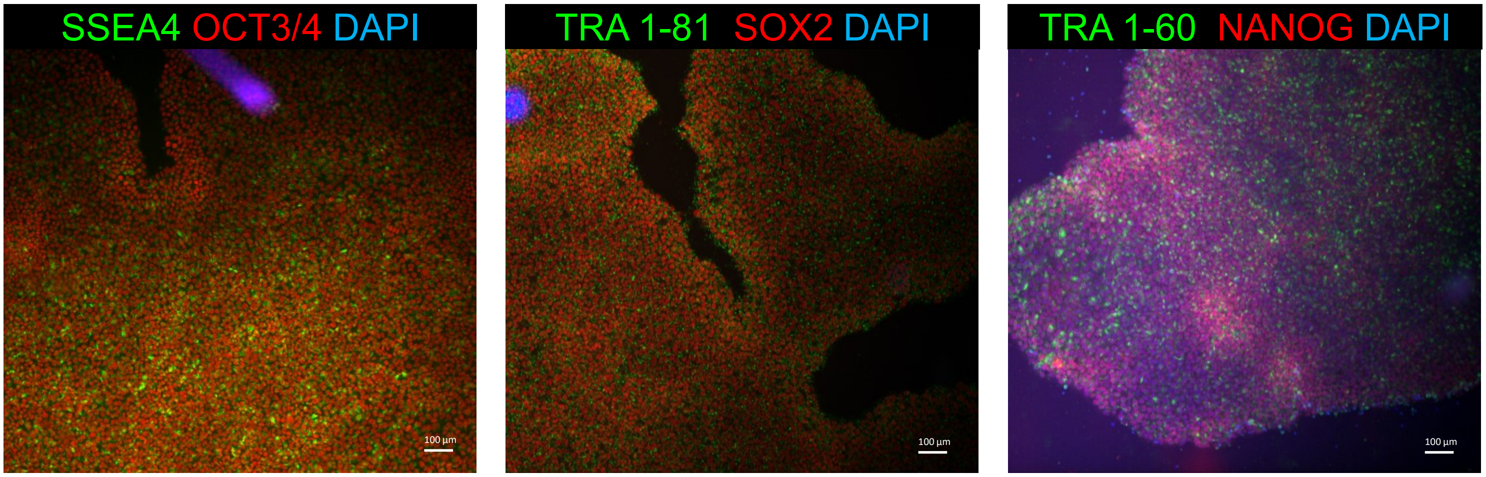


#### C. Pluritest

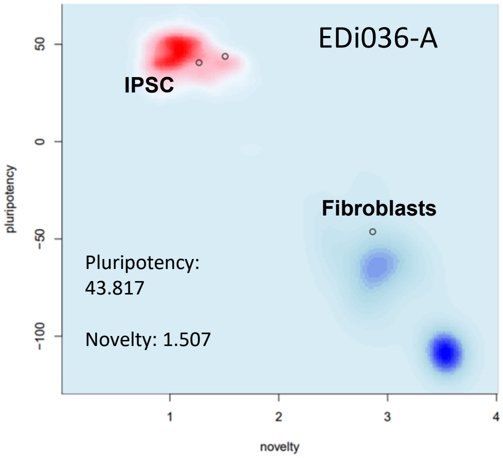


#### D. G -Band karyotype

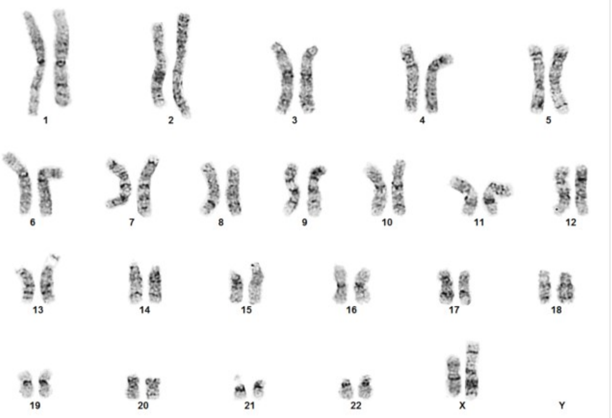


#### E. hPSC Scorecard

| IPSCs | | | | Embryoid Bodies | | | |
|---|---|---|---|---|---|---|---|
| Self-renew | Ecto | Meso | Endo | Self-renew | Ecto | Meso | Endo |
| + | − | − | − | − | + | + | + |
| 0.50 | 0.37 | 0.20 | -1.03 | -6.35 | 1.90 | 1.61 | 0.66 |

### sFig16: Characterization for iPSC line EDi037-A

## A. AP

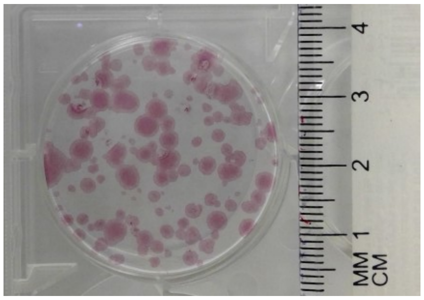


#### B. Immunocytochemistry

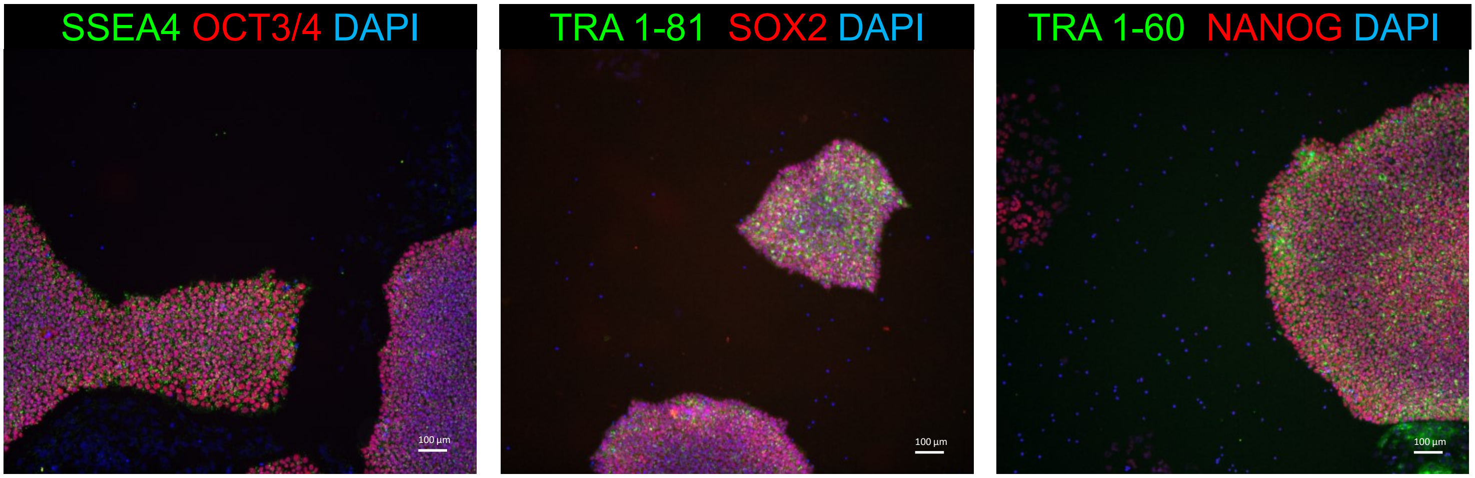


#### C. Pluritest

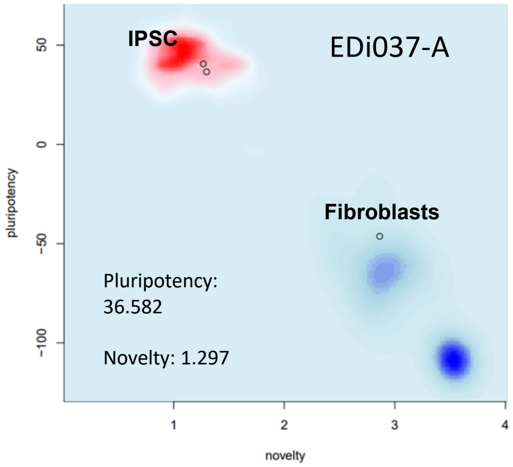


#### D. G -Band karyotype

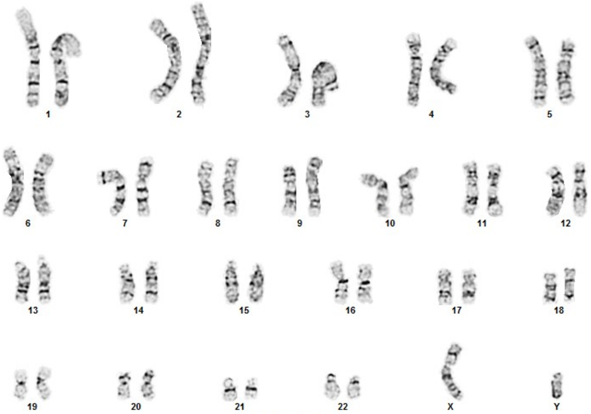


#### E. hPSC Scorecard

| IPSCs | | | | Embryoid Bodies | | | |
|---|---|---|---|---|---|---|---|
| Self-renew | Ecto | Meso | Endo | Self-renew | Ecto | Meso | Endo |
| + | − | − | − | − | + | + | + |
| 0.05 | 0.20 | 0.05 | -0.75 | -7.38 | 1.65 | 1.80 | 0.87 |

### sFig17: Characterization for iPSC line EDi038-A

## A. AP

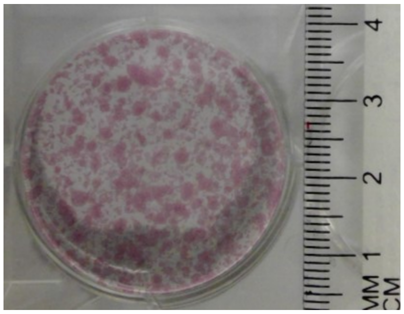

#### B. Immunocytochemistry

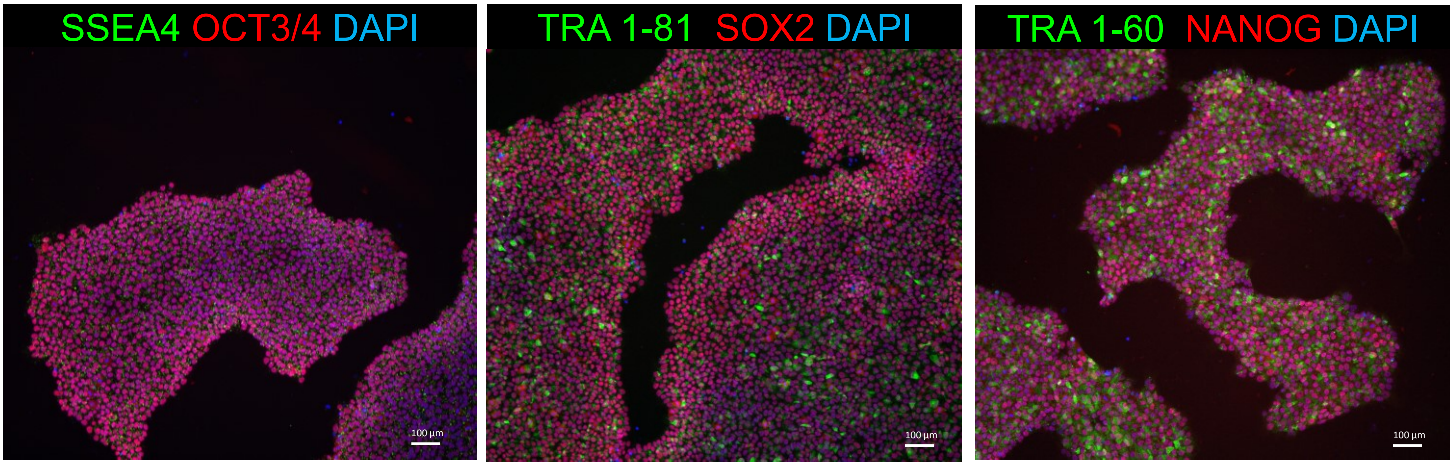


#### C. Pluritest

Plot unavailable

Pluripotency:
48.68024

Novelty:
1.418005

#### D. G-Band karyotype

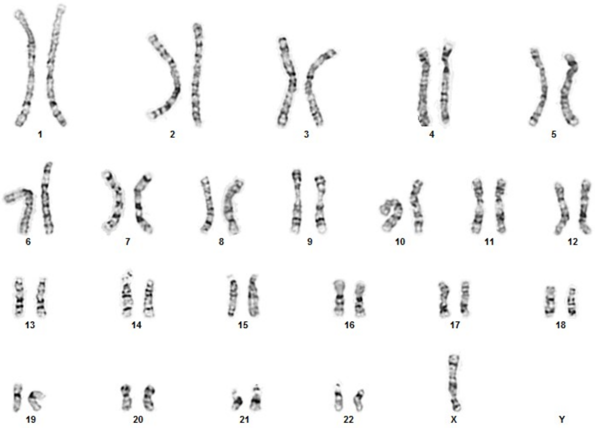

#### E. hPSC Scorecard

| IPSCs | | | | Embryoid Bodies | | | |
|---|---|---|---|---|---|---|---|
| Self-renew | Ecto | Meso | Endo | Self-renew | Ecto | Meso | Endo |
| + | − | − | − | − | + | + | + |
| 0.61 | -0.09 | 0.48 | -1.10 | -6.22 | 1.41 | 6.00 | 1.73 |

### sFig18: Characterization for iPSC line EDi039-A

## A. AP

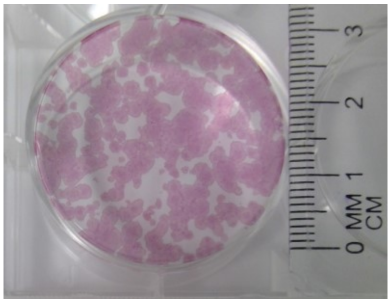


#### B. Immunocytochemistry

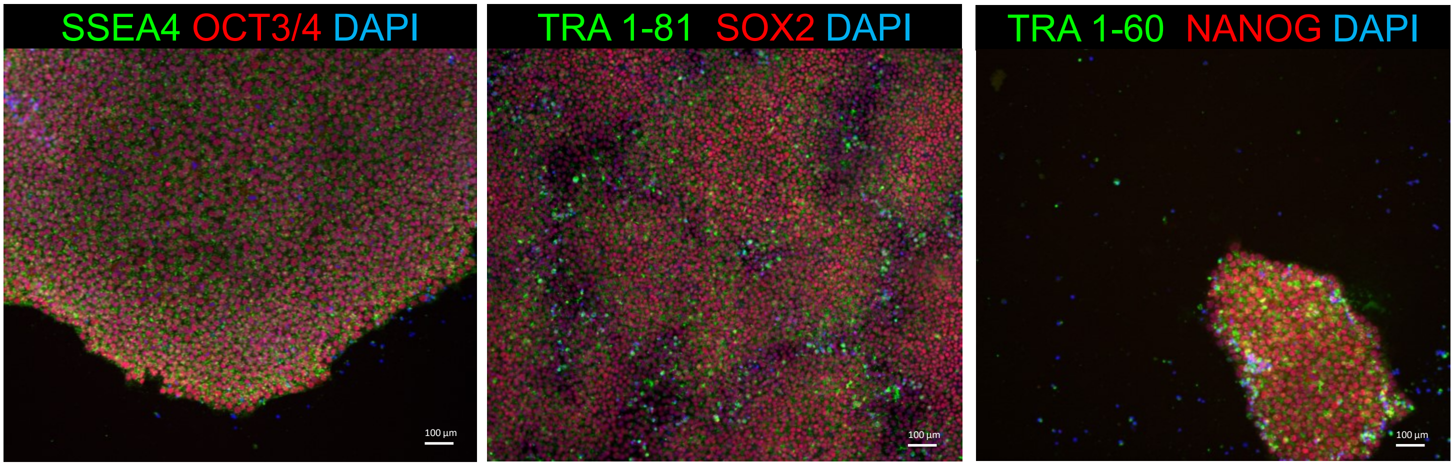


#### C. PluriTest

PluriTest not conducted.

#### D. G-Band karyotype

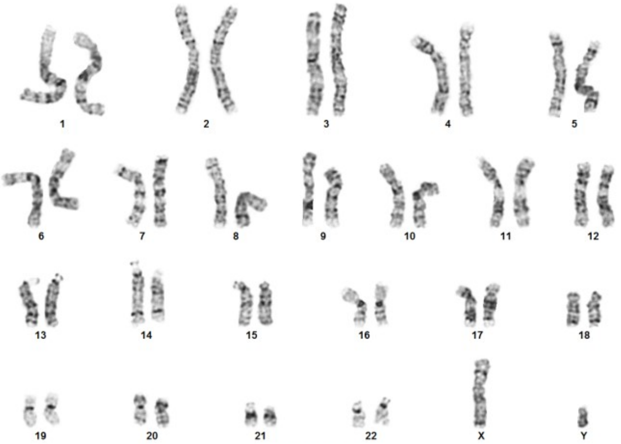


#### E. hPSC Scorecard

| iPSCs | | | | Embryoid Bodies | | | |
|---|---|---|---|---|---|---|---|
| Self-renew | Ecto | Meso | Endo | Self-renew | Ecto | Meso | Endo |
| + | − | − | − | − | + | + | + |
| -0.19 | 0.14 | -0.15 | -1.05 | -6.74 | 2.08 | 1.07 | 0.64 |

### sFig19: Characterization for iPSC line EDi040-A

## A. AP

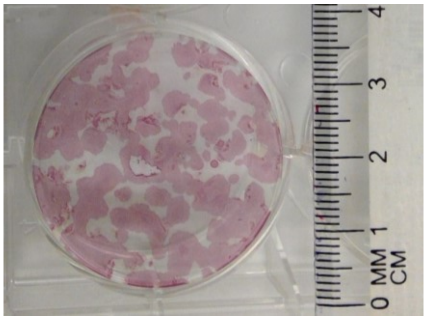


#### B. Immunocytochemistry

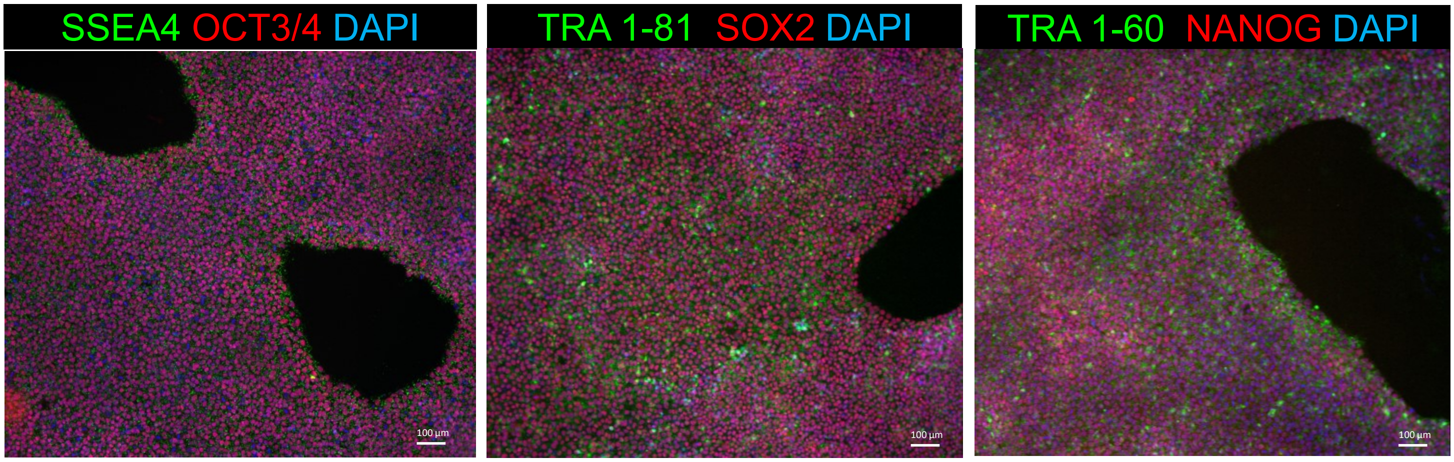


#### C. Pluritest

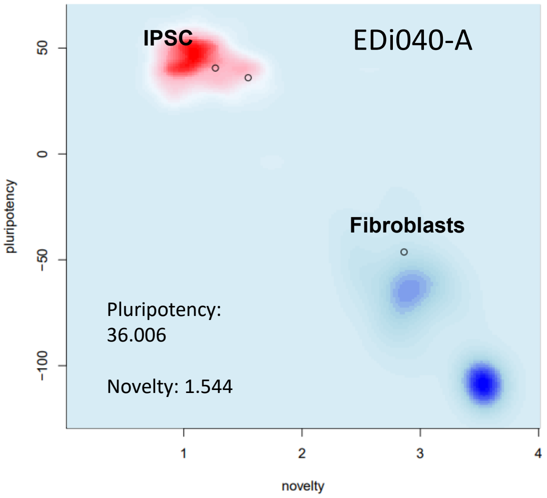


#### D. G-Band karyotype

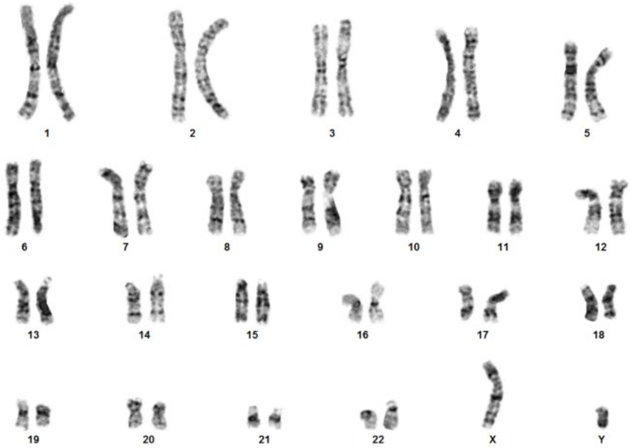


#### E. hPSC Scorecard

| IPSCs | | | | Embryoid Bodies | | | |
|---|---|---|---|---|---|---|---|
| Self-renew | Ecto | Meso | Endo | Self-renew | Ecto | Meso | Endo |
| + | − | − | − | − | + | + | + |
| 0.01 | -0.45 | 0.24 | -0.94 | -3.18 | 1.57 | 0.98 | -0.56 |

### sFig20: Characterization for iPSC line EDi041-A

## A. AP

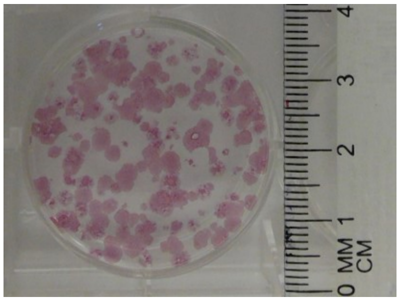


#### B. Immunocytochemistry

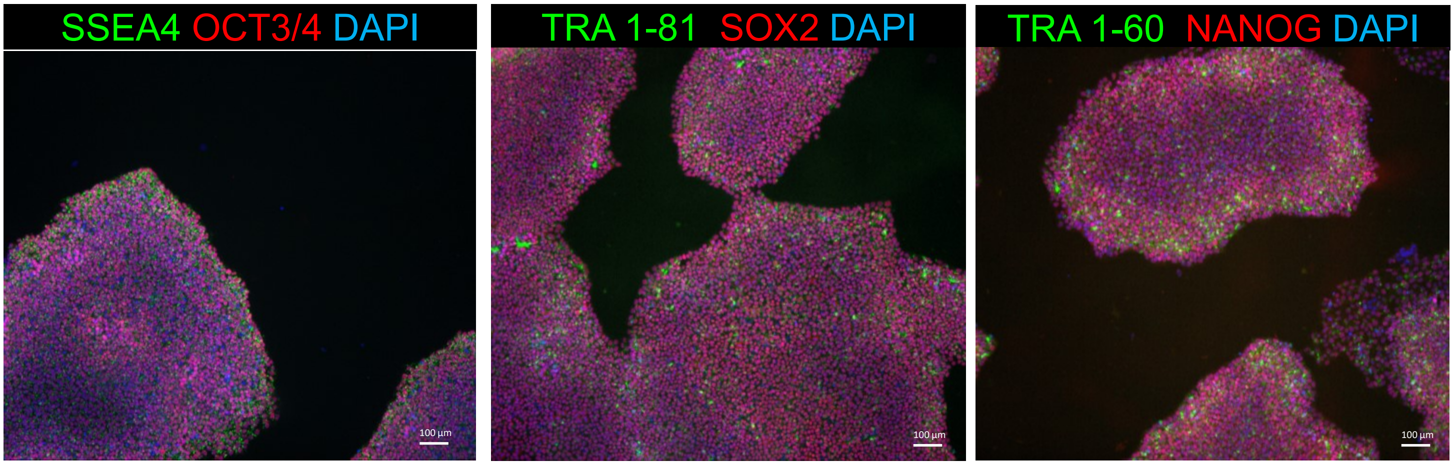


#### C. Pluritest

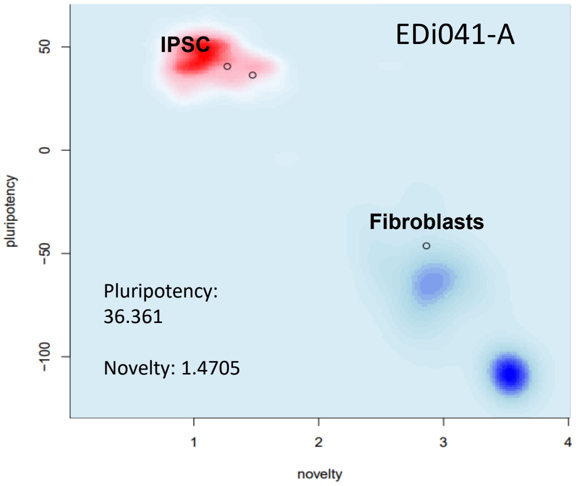


#### D. G-Band karyotype

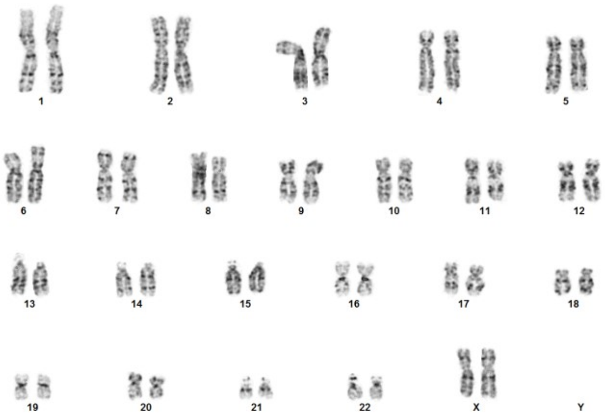


#### E. hPSC Scorecard

| iPSCs | | | | Embryoid Bodies | | | |
|---|---|---|---|---|---|---|---|
| Self-renew | Ecto | Meso | Endo | Self-renew | Ecto | Meso | Endo |
| + | − | − | − | − | + | + | + |
| --0.18 | 0.05 | -0.32 | -1.24 | -7.04 | 1.61 | 1.42 | 0.67 |

### sFig21: Characterization for iPSC line EDi042-A

## A. AP

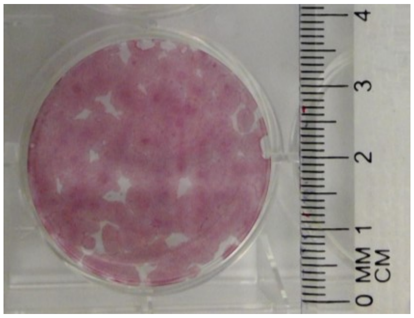


#### B. Immunocytochemistry

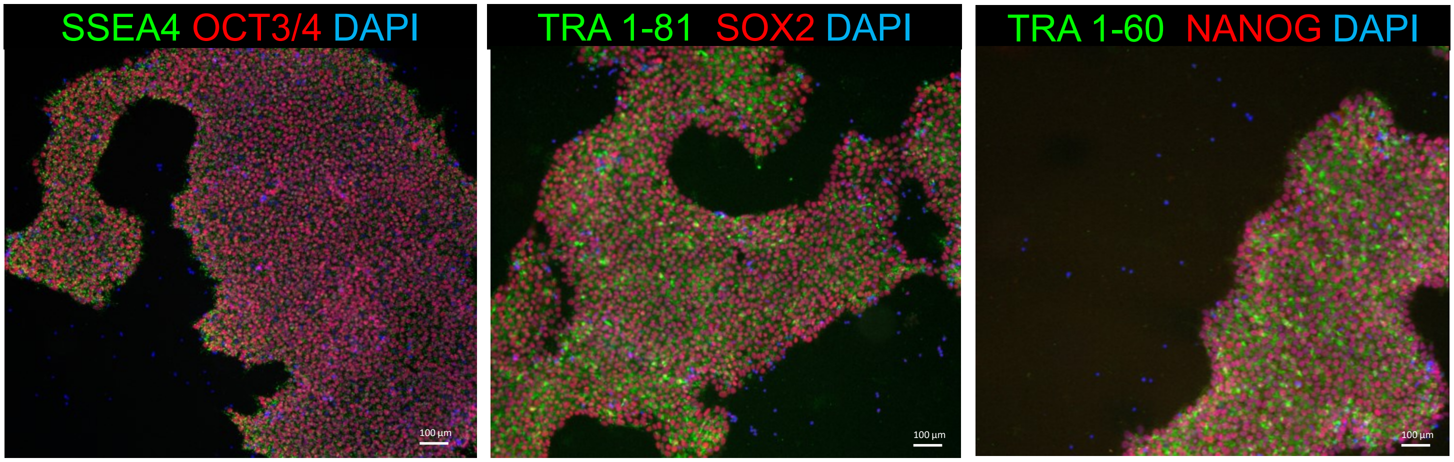


#### C. Pluritest

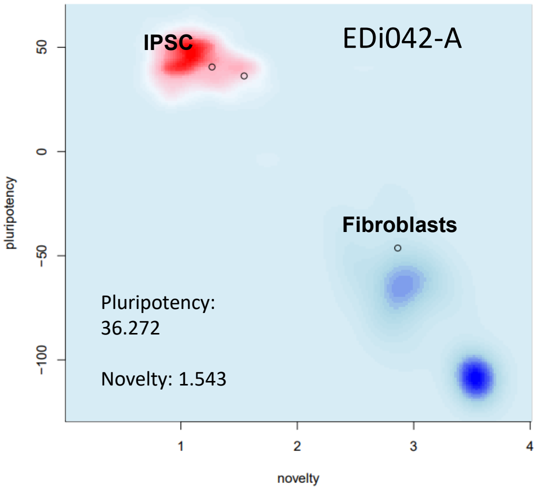


#### D. G-Band karyotype

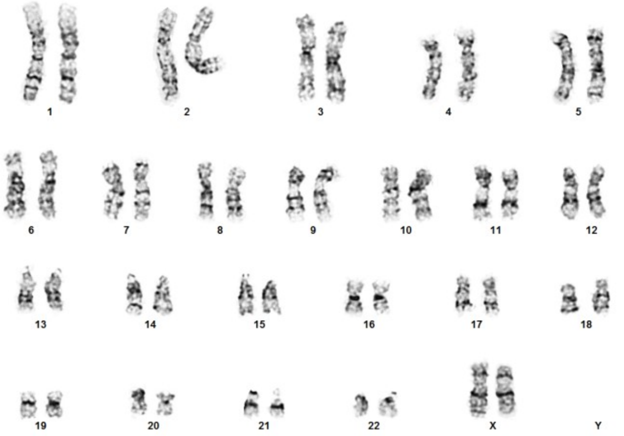


#### E. hPSC Scorecard

| IPSCs | | | | Embryoid Bodies | | | |
|---|---|---|---|---|---|---|---|
| Self-renew | Ecto | Meso | Endo | Self-renew | Ecto | Meso | Endo |
| + | − | − | − | − | + | + | − |
| 0.05 | -0.13 | 0.05 | -1.04 | -5.15 | 2.62 | 1.07 | 0.22 |

### sFig22: Characterization for iPSC line EDi043-A

## A. AP

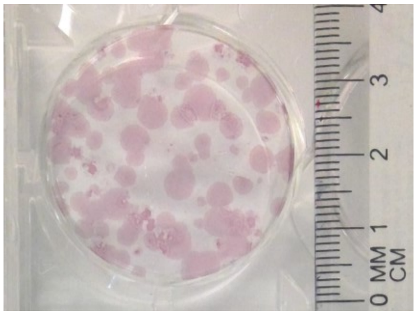


#### B. Immunocytochemistry

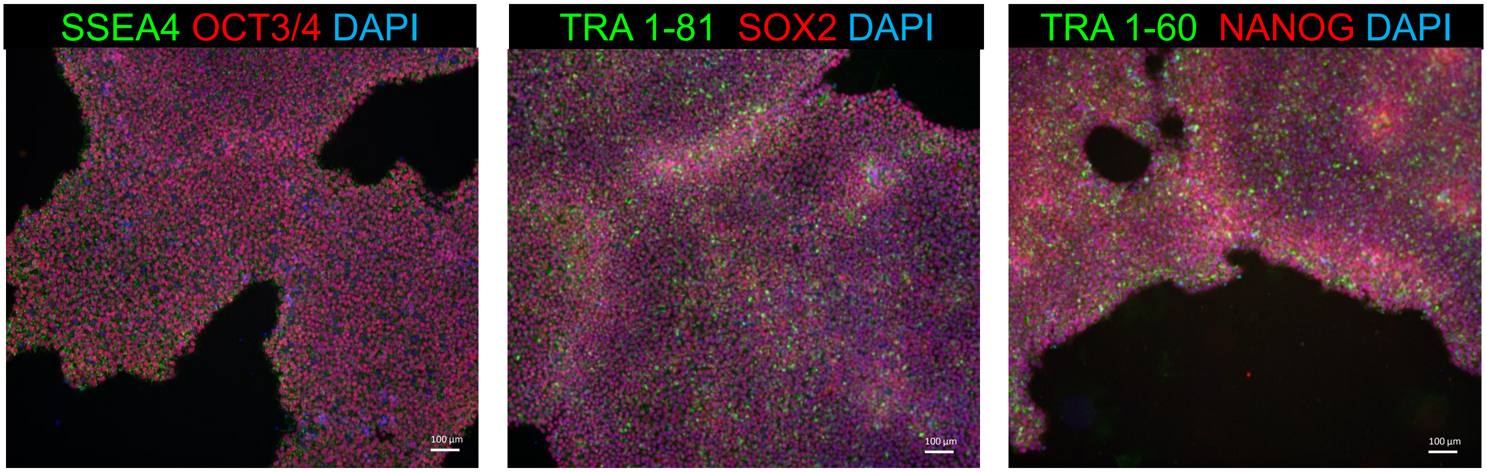


#### C. Pluritest

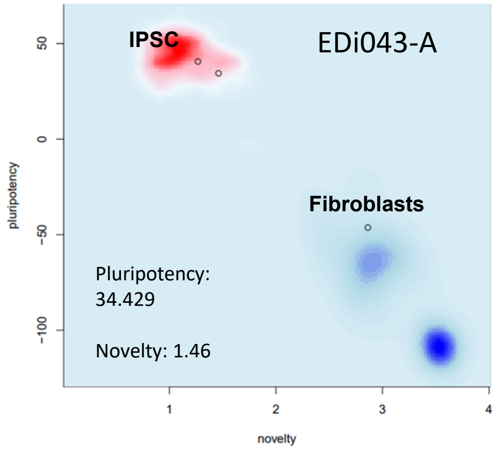


#### D. G -Band karyotype

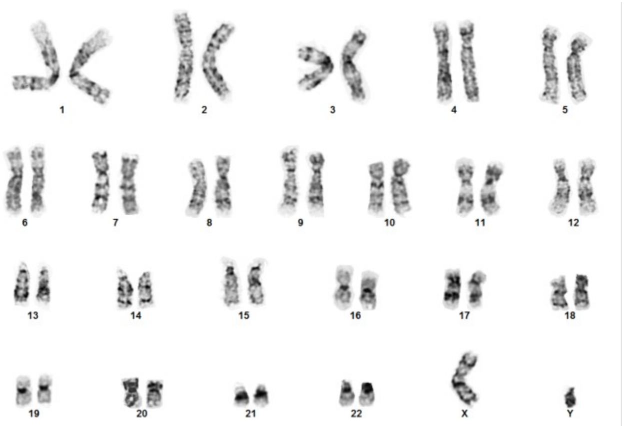


#### E. hPSC Scorecard

| IPSCs | | | | Embryoid Bodies | | | |
|---|---|---|---|---|---|---|---|
| Self-renew | Ecto | Meso | Endo | Self-renew | Ecto | Meso | Endo |
| + | − | − | − | − | + | + | + |
| -0.03 | 0.38 | 0.07 | -1.23 | -6.46 | 1.59 | 2.57 | 1.19 |

### sFig23: Characterization for iPSC line EDi044-A

## A. AP

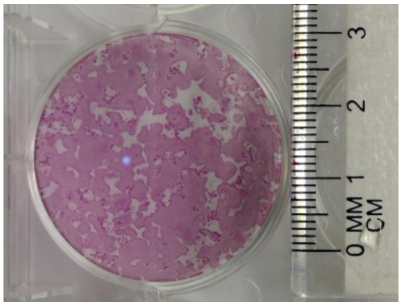

#### B. Immunocytochemistry

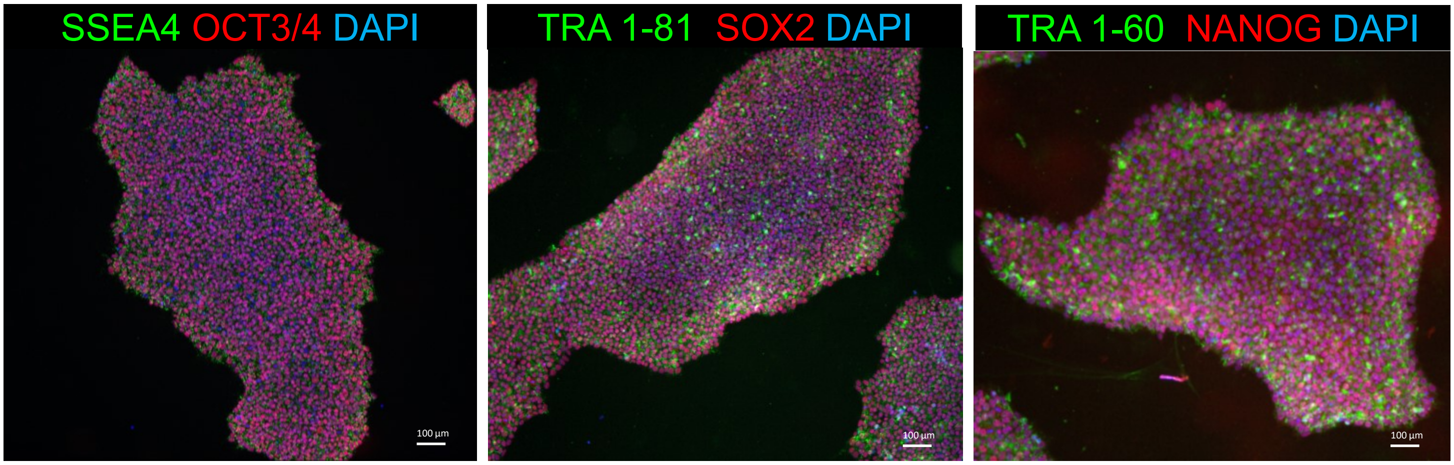


#### C. Pluritest

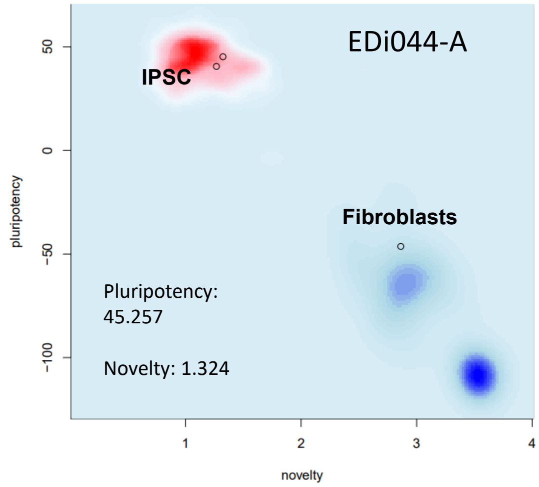


#### D. G-Band karyotype

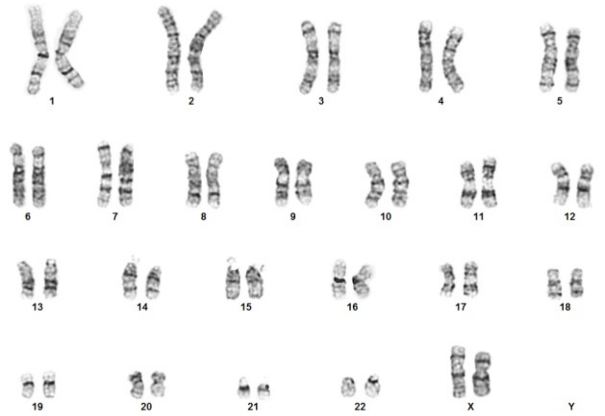

#### E. hPSC Scorecard

| iPSCs | | | | Embryoid Bodies | | | |
|---|---|---|---|---|---|---|---|
| Self-renew | Ecto | Meso | Endo | Self-renew | Ecto | Meso | Endo |
| -0.75 | -0.19 | 0.58 | -0.80 | -6.29 | 1.65 | 3.42 | 0.94 |

### sFig24: Characterization for iPSC line EDi045-A

## A. AP

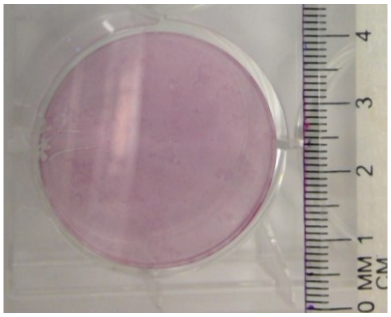


#### B. Immunocytochemistry

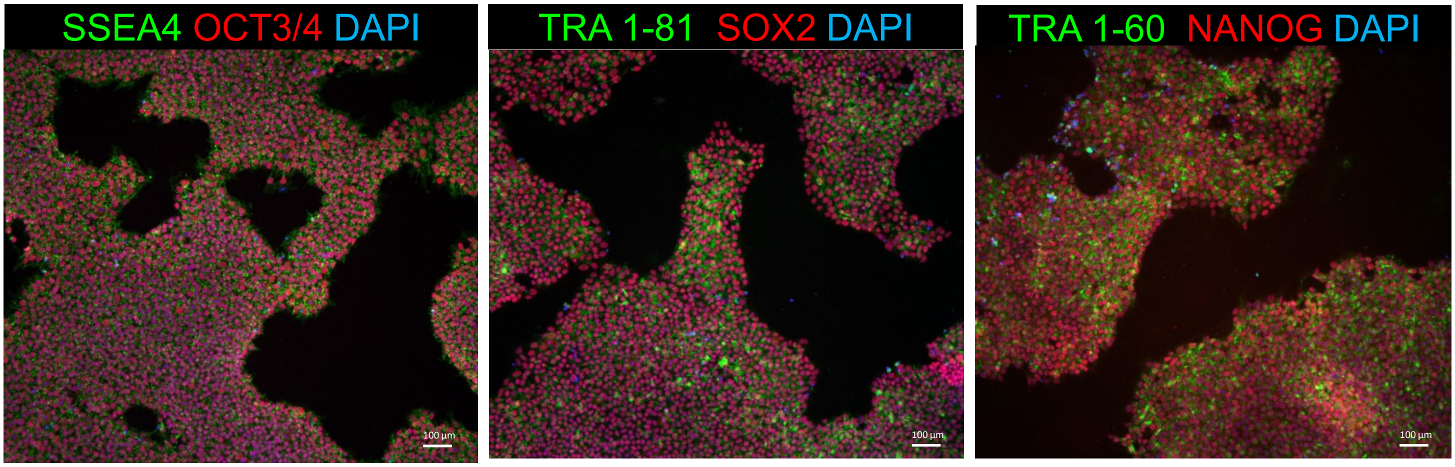


#### C. Pluritest

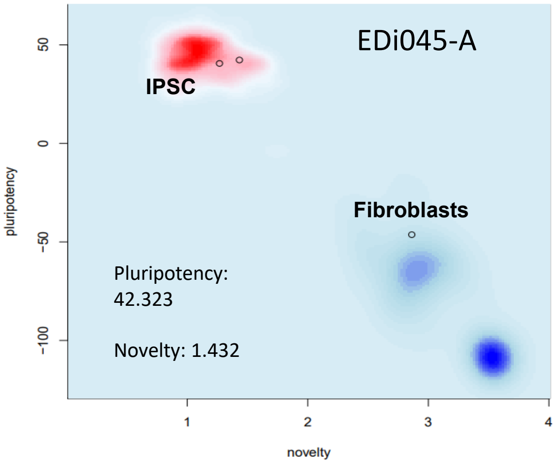


#### D. G-Band karyotype

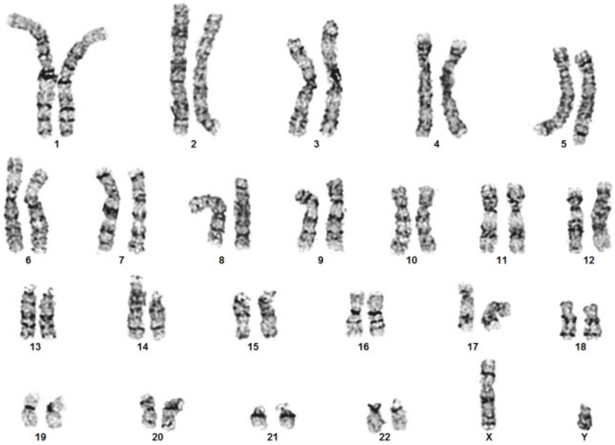


#### E. hPSC Scorecard

| IPSCs | | | | Embryoid Bodies | | | |
|---|---|---|---|---|---|---|---|
| Self-renew | Ecto | Meso | Endo | Self-renew | Ecto | Meso | Endo |
| + | − | − | − | − | + | + | + |
| -0.11 | 0.15 | 0.22 | -0.81 | -6.43 | 2.29 | 1.53 | 0.62 |
